# Spatial-ID: a cell typing method for spatially resolved transcriptomics via transfer learning and spatial embedding

**DOI:** 10.1101/2022.05.26.493527

**Authors:** Rongbo Shen, Lin Liu, Zihan Wu, Ying Zhang, Zhiyuan Yuan, Junfu Guo, Fan Yang, Chao Zhang, Bichao Chen, Chao Liu, Jing Guo, Guozhen Fan, Yong Zhang, Yuxiang Li, Xun Xu, Jianhua Yao

**Affiliations:** Tencent AI Lab, Shenzhen, China; BGI-Shenzhen, Shenzhen 518083, China; Guangdong Bigdata Engineering Technology Research Center for Life Sciences, Shenzhen 518083, China; MOE Key Laboratory of Bioinformatics; Bioinformatics Division and Center for Synthetic & Systems Biology, BNRist; Department of Automation, Tsinghua University, Beijing 100084, China

## Abstract

Spatially resolved transcriptomics (SRT) provides the opportunity to investigate the gene expression profiles and the spatial context of cells in naive state. Cell type annotation is a crucial task in the spatial transcriptome analysis of cell and tissue biology. In this study, we propose Spatial-ID, a supervision-based cell typing method, for high-throughput cell-level SRT datasets that integrates transfer learning and spatial embedding. Spatial-ID effectively incorporates the existing knowledge of reference scRNA-seq datasets and the spatial information of SRT datasets. A series of quantitative comparison experiments on public available SRT datasets demonstrate the superiority of Spatial-ID compared with other state-of-the-art methods. Besides, the application of Spatial-ID on a SRT dataset with 3D spatial dimension measured by Stereo-seq shows its advancement on the large field tissues with subcellular spatial resolution.

## Introduction

In the last decade, single-cell RNA sequencing (scRNA-seq) technologies have made significant progress towards the systematic characterization of cell dynamics^1,2^. However, the dissociation step of scRNA-seq leads to loss of spatial information, preventing the investigation of tissue organization in its naive state^3^. To reveal fine-scale spatial organization and microenvironments within tissues, various spatially resolved transcriptomics (SRT)^4–6^ technologies have been introduced to map gene expression profiles to the spatial context. Early technologies attempted to spatially map multiplexed gene expression by either in situ hybridization (ISH)^7,8^ or spatially barcoded oligo-deoxythymidine microarrays^9,10^. More advanced ones, e.g., seqFISH^11^, seqFISH+^12^, MERFISH^13^, Slide-seq^14,15^, HDST^16^ and Stereo-seq^17^, have been proposed to improve in terms of spatial resolution, molecular throughput and transcript detection sensitivity, allowing for comprehensive mapping of transcripts over large tissue sections.

Cell type annotation is a fundamental task for cell and tissue biology^70^ that can help characterize the biological process of tissues at single cell level. This task is conventionally performed by single cell transcriptome analysis on the data acquired by scRNA-seq technology. Facing the exponential growth of high-dimensional and noisy data associated with sequencing technologies^18^, it requires high performance annotation methods that can effectively reduce dimensionality and are robust to data noise^19^. Annotating cell identities is even more challenging for SRT datasets^20^. For example, spot-based protocols such as Visium^9^, Slide-seq^14,15^, HDST^16^ and Stereo-seq^17^ capture RNA from areas spanning more (or less) than one cell, without consideration of cell boundaries. In addition, due to inherited experimental design, the low transcript detection sensitivity of spot-based protocol further makes the measured transcriptomic profiles deviated from the real transcript levels of single cells. Also, FISH-based protocols face similar problem arising from potential inaccuracy of cell segmentation.

To completely understand the cell type organization of biological tissues, many cell atlas projects such as Human Cell Atlas (HCA)^21^, Allen Brain Atlas (ABA)^22^, Brain Atlas of BICCN^23^, exploited hierarchical clustering scheme on large-scale scRNA-seq datasets to establish cell type taxonomy and define marker genes of each cell type by differential gene expression analysis. One of the main limitations of clustering methods is that the cell type taxonomy and marker gene sets are bound to a certain level of clustering resolution^24^. Moreover, the computational characterization of the cell heterogeneity requires a series of statistical per-cell gene signature assessments to remove low-quality and doublet-driven clusters^3,24^. Besides, facing the explosive data growth of new sequencing technologies, such de novo clustering methods^24^ become more laborious, computationally intensive and inefficient. It is more desirable to develop supervised cell typing methods that transfer information from annotated reference datasets to newly generated datasets^75^.

To better characterize cell identities from reference datasets with well-defined cell types, many correlation-based and supervision-based cell typing methods have been introduced in scRNA-seq data analysis, such as Seruat v3^25^, SingleR^26^, Scmap^27^, Cell-ID^28^, ScNym^29^, SciBet^30^ and ScDeepSort^31^. Seruat v3^25^ provided a unified cell typing strategy to transfer information from a reference dataset to newly sequenced dataset, which could integrate diverse single-cell datasets across different single-cell sequencing technologies and omics. SingleR^26^ performed cell type annotation for newly sequenced single-cell transcriptomics dataset that was correlated to reference datasets of pure cell types and strengthened its inferences by reducing the reference datasets to only top cell types iteratively. Scmap^27^ provided a cell typing strategy that projected a newly sequenced cell onto a reference dataset by searching the most similar clusters or cells (i.e., nearest neighbors) in the reference dataset and then assigning a cell type if its nearest neighbors have the same cell type. Cell-ID^28^ independently extracted per-cell gene signatures for a newly sequencing dataset and reference datasets through multiple correspondence analysis, then used per-cell gene signatures to perform automatic cell type and functional annotation for target single-cell transcriptomic dataset by cell matching and label transferring from reference datasets. ScNym^29^ employed an adversarial neural network to transfer cell identity annotations from a labeled reference dataset to an unlabeled newly sequencing dataset despite biological and technical differences. SciBet^30^ used the mean expression of cell type-specific genes selected by E-test to train a multinomial-distribution model, then calculated the likelihood function of a test cell using the trained model and annotated cell type for the test cell with maximum likelihood estimation. ScDeepSort^31^ pre-trained a weighted graph neural network (GNN)^38^ to perform cell type annotation for newly sequencing single cell transcriptomic datasets. However, applying these correlation-based and supervision-based cell typing methods to SRT datasets does not efficiently use the available spatial information which may be beneficial to cell type annotation.

By considering the characteristics of the SRT technologies, several SRT analysis methods such as deconvolution-based, PGM-based^34^ and spatial embedding-based were proposed. The deconvolution-based methods^32,33^ learned cell-type gene signatures from reference datasets to decompose sequenced spots in SRT datasets, where a spot contained multiple cells. However, the deconvolution-based methods are not suitable for the recent SRT technologies with cell-level spatial resolution. BayesSpace^34^, a PGM-based SRT analysis method, employed a Markov random field (MRF) to perform clustering analysis and resolution enhancement for spatial domain analysis of SRT datasets. Several spatial embedding-based clustering methods such as SpaGCN^34^, SEDR^36^, STAGATE^37^ were proposed to spatial domain clustering analysis that enable coherent gene expression in spatial domains by integrating gene expression and spatial location together. Inspired by the significant advancements of these spatial embedding-based clustering methods in identifying anatomical spatial domains, embedding spatial information should be beneficial to cell type annotation of SRT datasets.

In this study, we propose a cell typing method (SPATIAL cell type IDentification, Spatial-ID) that integrates transfer learning and spatial embedding strategies for high-throughput cell-level SRT datasets. The transfer learning strategy employs scRNA-seq datasets with well-defined cell-type gene signatures collected from similar tissues to train deep neural network (DNN) models. The cell type taxonomy can also be aligned with existing cell atlas that was constructed from similar tissues. To perform spatial embedding, we propose a graph convolution network (GCN)^38^ that constructs a spatial neighbor graph by considering each cell as a node and the spatial location relationships between cells as edges. In the architecture of GCN, an autoencoder^39^ is used to encode gene expression and a variational graph autoencoder^40^ is used to embed spatial information simultaneously. To handle the large number of cells in the high-throughput SRT data, we employ sparse convolution in GCN to accelerate the framework^41^. A self-supervised learning strategy is performed by constructing pseudo-labels from the probability distribution predicted by the DNN model of transfer learning^42^.

The main contribution of the Spatial-ID is the effective incorporation of existing knowledge of reference scRNA-seq datasets and the spatial information of SRT datasets. A series of comparison experiments on public available SRT datasets with different data characteristics (See Table 1) demonstrate the superiority of Spatial-ID in cell type annotation compared with other state-of-the-art methods (See Table 2), i.e., Seruat v3, SingleR, Scmap, Cell-ID, ScNym and SciBet. Furthermore, Spatial-ID can effectively perform cell type annotation for 3D SRT datasets. Moreover, the extended experiments of new cell type discovery demonstrate that Spatial-ID has a promising ability to detect new cell types. A group of simulation experiments with different gene dropout rates demonstrates more robustness of Spatial-ID than other state-of-the-art methods. These results suggest that embedding spatial information can substantially improve cell type annotation on SRT datasets. Besides, the application of Spatial-ID on a Stereo-seq SRT dataset with 3D spatial dimension shows its advancement on the large field tissues (∼1cm^2^) with subcellular spatial resolution. The cell types identified by Spatial-ID present high consistency with previous studies. Besides, by mapping the identified cell types with identified spatial gene patterns, the significant GO terms of the spatial gene patterns further reveal the functions and underlying biological processes of the identified cell types.

**Table 1.** Datasets used in this study.

| SRT datasets/ Tissue sample | Techniques | Characteristics |  |  |  |  | Reference scRNA-seq datasets |  |
| --- | --- | --- | --- | --- | --- | --- | --- | --- |
|  |  | Samples | Cells | Genes | Field size | Notes | Name | meta information |
| Mouse brain - primary motor cortex (MOP), (https://doi.org/10.35077/g.21) | High-plex RNA imaging - Multiplexed error-robust FISH (MERFISH) | 12 | 280,186 | 254 | ~2.0mm×2.4mm | Samples are collected from 2 mouse brains, where each mouse contains 6 samples, and each sample includes 4-6 coronal slices (10um thick) that are imaged together on the same coverslip. | snRNA-seq 10x v3 B, (https://assets.nemoarc-hive.org/dat-ch1nqb7) | 159,738 cells, 31,053 genes. |
| Mouse brain - hypothalamic preoptic region (3D), (https://datadryad.org/stash/dataset/doi:10.5061/dryad.8t8s248) | High-plex RNA imaging - Multiplexed error-robust FISH (MERFISH) | 3 | 213,192 | 155 | 1.8mm×1.8mm×0.6mm | 3 samples with naive behavior from 2 female mice and 1 male mouse brains (Bregma 0.26 to -0.29), each sample includes 12 slices with 50um interval. | GSE113576, (https://www.ncbi.nlm.nih.gov/geo/query/acc.cgi?acc=GSE113576) | 31,299 cells, 27,998 genes. |
| Mouse spermatogenesis, (https://www.dropbox.com/s/ygzpj0d0oh67br0/Testis_Slideseq_Data.zip?dl=0) | In situ capture, Spatial barcoding - Slide-seq, 10um diameter per sequencing spot. | 6 | 207,335 | 27,181 | ~2.5mm×2.5mm | 3 samples from leptin-deficient diabetic mice (ob/ob) and 3 samples from wild-type (WT) mice. | GSE112393, (https://www.ncbi.nlm.nih.gov/geo/query/acc.cgi?acc=GSE112393) | 34,633 cells, 37,241 genes. |
| Mouse brain hemisphere (3D) | In situ capture, Spatial barcoding - Spatio-Temporal Enhanced REsolution Omics-sequencing (Stereo-seq), 0.22um diameter per sequencing spot and 0.5um center-to-center distance. | 3 | 177,077 | 26,219 | ~1cm×1cm×0.02mm | 3 adjacent coronal sections (10μm thick, without intervals) along the anterior-posterior axis (Bregma -3.56 to -3.66) from a 5-week-old C57BL/6J male mouse brain. | Mouse brain atlas of cell types from the Linnarsson Lab, (http://mousebrain.org/adolescent/) | 160,796 cells, 27,998 genes. |

**Table 2.** The control methods. The supervision-based methods employ reference scRNA-seq datasets as trainset to train models. The robustness indicates the variations of performance across different datasets. The usability indicates the degree of easy-to-use. The efficiency indicates the running efficiency.

| Algorithm | Paradigm | Robustness | Usability | Efficiency | Languages | Mechanism notes | Strategies |
| --- | --- | --- | --- | --- | --- | --- | --- |
| Seruat v3 | Correlation-based | ✓ | ✓ | – | R | Seruat v3 transfers information from a reference dataset to newly sequencing dataset, which could integrate diverse single-cell datasets across different single-cell sequencing technologies and omics datasets | 1. Integrating diverse single-cell datasets across technologies and modalities.<br>2. Projecting the datasets into a subspace defined by shared correlation structure across datasets.<br>3. Using canonical correlation analysis (CCA) and mutual nearest neighbors (MNNs) to identify shared subpopulations across datasets. |
| SingleR | Correlation-based | ✓ | ✓ | – | R | SingleR correlates single-cell transcriptomes with reference transcriptomic datasets and improve its inferences by reducing the reference datasets to only top cell types iteratively. | 1. Identifying variable genes among cell types in the reference dataset.<br>2. Correlating each single-cell transcriptome with each sample in the reference dataset using Spearman coefficient.<br>3. Reducing the reference dataset to only top cell types by iterative fine-tuning. |
| Scmap | Correlation-based | – | – | – | R | Scmap projects a newly sequenced cell onto a reference dataset by searching the most similar clusters or cells (i.e., nearest neighbors) in the reference dataset and then assigning a cell type if its nearest neighbors have the same cell type. | 1. A method conceptually similar to M3Drop is used to select informative genes, which can overcome batch effects.<br>2. Approximate nearest neighbor search using product quantizer.<br>3. The similarity value are measured by Pearson, Spearman and cosine. |
| Cell-ID | Correlation-based | – | ✓ | – | R | Cell-ID extracts per-cell gene signatures for a newly sequencing dataset and reference datasets through multiple correspondence analysis (MCA), then uses per-cell gene signatures to perform automatic cell type and functional annotation for target single-cell transcriptomic dataset by cell matching and label transferring from reference datasets. | 1. A statistical technique (i.e., MCA) provides a simultaneous representation of cells and genes in low-dimensional space.<br>2. A gene ranking is obtained for each cell in a dataset based on their distance in the MCA space, where the top-ranked genes for a given cell define its gene signature. |
| ScNym | Supervision-based | ✓ | – | ✓ | Python | ScNym employs an adversarial neural network to transfer cell identity annotations from a labeled reference dataset to an unlabeled newly sequencing dataset despite biological and technical differences. | 1. A semisupervised adversarial neural network is employed to eliminate domain shifts across different datasets.<br>2. Training and target cell profiles and their labels are augmented using weighted averages. |
| SciBet | Supervision-based | – | ✓ | ✓ | R | SciBet uses the mean expression of cell type-specific genes to train a multinomial-distribution model, then calculated the likelihood function of a test cell using the trained model and annotated cell type for the test cell with maximum likelihood estimation. | 1. The cell type-specific genes are selected from the training set by E-test.<br>2. The cell type is assigned by the highest likelihood achieved by the model. |

## Results

### The pipeline of Spatial-ID

The pipeline of our proposed Spatial-ID is shown in Fig. 1. Given a SRT dataset, the DNN model of transfer learning generates the probability distribution of each cell, which is used to construct pseudo-label (Fig. 1b) through a temperature setting strategy of knowledge transfer (See Stage 1 and Methods). Meanwhile, the autoencoder module (Fig. 1c) learns encoded gene representations from the gene expression matrix, and a spatial neighbor graph is constructed through calculating the Euclidean distances between cells (Fig. 1d). Next, the variational graph autoencoder incorporates the encoded gene representations and the spatial neighbor graph to produce spatial embeddings (Fig. 1e). Then, the encoded gene representations and spatial embeddings are aggregated to generate the final latent representations that provide comprehensive characters of gene expression and spatial information. Thereafter, the final latent representations are used to reconstruct the gene expression in the autoencoder and the spatial neighbor graph in the variational graph autoencoder. Simultaneously, using the generated pseudo-labels, the self-supervised learning strategy employs final latent representations to train the classifier (See Stage 2). After training, our framework generates the final class probability distribution of each cell in a given SRT dataset (See Stage 3).

**Fig. 1.**
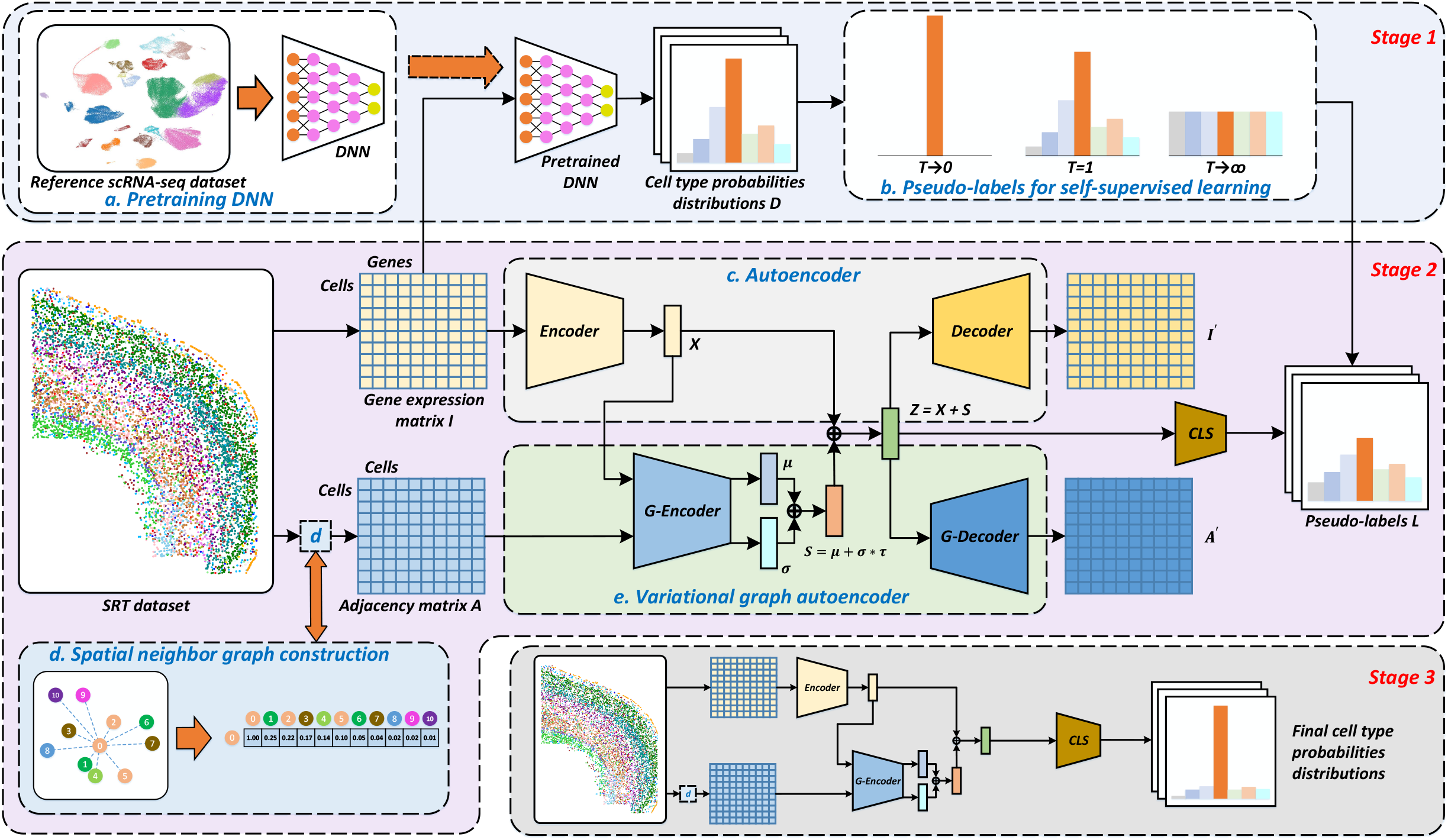
Overview of Spatial-ID. Stage1 involves knowledge transfer from reference datasets. Stage 2 involves feature embedding of gene expression and spatial information, and classifier learning. Stage 3 uses the model trained in Stage 2 to perform cell type annotation. **(a)** Reference scRNA-seq datasets are employed to pretrain deep neural network (DNN) models. (**b)** Based on the cell type probabilities distributions *D* produced by pretrained DNN, pseudo-labels *L* are generated by adjusting the temperature parameter *T*. (**c)** A deep autoencoder is used to learn encoded gene representation *X* through reducing the dimension of the gene expression matrix *I*. The gene expression matrix *I*′ reconstructed by decoder is used to optimize the autoencoder by minimizing with the input gene expression matrix *I*. (**d)** A spatial neighbor graph is constructed to represent the spatial relationships between neighboring cells, where the relationship weight of each pair of cells is negatively associated with Euclidean distance. Therefore, the spatial neighbor graph is represented as an adjacency matrix *A*. (**e)** A variational graph autoencoder (VGAE, a kind of GCN) is used to embed the encoded gene representations *X* from autoencoder and the adjacency matrix *A*, and then generate the spatial embedding *s* as output. The reconstructed adjacency matrix *A^′^* is used to optimize the VGAE by minimizing with the input adjacency matrix *A*.

To verify the advancement of the proposed Spatial-ID, we apply Spatial-ID on 4 representative SRT datasets with different characteristics (See Table 1), including two public available FISH-based SRT datasets (i.e., mouse primary motor cortex^43^ and mouse hypothalamic preoptic region^44^) generated by MERFISH, a public available Slide-seq SRT dataset (i.e., mouse spermatogenesis^45^), and a self-collected Stereo-seq SRT dataset (i.e., large field mouse brain hemisphere). The two FISH-based datasets measure hundreds of genes, in which the mouse hypothalamic preoptic region dataset is a 3D SRT dataset that provides continuous slices with equal interval. The spot-based Slide-seq SRT dataset and the Stereo-seq SRT dataset measure tens of thousands of genes, and the last one provides 3D spatial dimension, subcellular spatial resolution and large field view on entire mouse brain hemisphere. The two FISH-based SRT datasets and the Slide-seq SRT dataset are employed to quantitatively compare Spatial-ID with other state-of-the-art methods (See Table 2). The Stereo-seq SRT dataset is employed to demonstrate the advancement of Spatial-ID on spatial transcriptomic analysis of the large field tissues. Besides, to demonstrate the adaptation and robustness of Spatial-ID, an extend experiment of new cell type discovery and a simulation experiment of different gene dropout rate are performed on the FISH-based mouse primary motor cortex dataset.

### Application to mouse primary motor cortex dataset measured by MERFISH

We first perform a quantitative comparison between the Spatial-ID and the control methods on the mouse primary motor cortex (MOP, See Fig. 2a) dataset^43^ measured by MERFISH. The MOP SRT dataset contains 12 samples, including total 280,186 cells and 254 genes. The snRNA-seq 10x v3 B dataset^46^ is used as the trainset of the DNN model in Spatial-ID and the reference dataset in the control methods, which contains 159,738 cells and 31,053 genes. The MOP SRT dataset and the snRNA-seq 10x v3 B dataset are derived from Brain Atlas of BICCN^23^, thus the cell type assignments of them adapt the same MOP cell taxonomy that is a hierarchical organization with reference to the common cell type nomenclature (CCN)^47^ found by Allen Institute. The cells are divided into excitatory neuronal cells (glutamatergic), inhibitory neuronal cells (GABAergic) and non-neuronal cell classes at the first level, then cells in each class are further divided into more subclasses based on their marker genes or spatial organization in cortex (Fig. 2b, Fig. 2d and Extend Fig. 1a). For example, the inhibitory neurons are further divided into five subclasses by marker genes: parvalbumin (Pvalb), somatostatin (Sst), vasoactive intestinal polypeptide (Vip), synuclein gamma (Sncg) and lysosomal-associated membrane protein family member 5 (Lamp5) (Extend Fig. 1c). The excitatory neurons are further divided into several layers with distinct projection properties (defined by known marker genes): intra-telencephalic neurons (L2/3 IT, L4/5 IT, L5 IT, L6 IT) (Extend Fig. 1b), extra-telencephalic projecting neurons (L5 ET), near-projecting neurons (L5/6 NP), corticothalamic projection neurons (L6 CT) and layer 6b neurons (L6b) (Fig. 2h). It should be noted that the number of cell types, the number of measured genes, relative abundance of cells in various cell types and the gene dropout rate differ in the snRNA-seq 10x v3 B dataset and the MOP ST dataset. For example, smooth muscle cells (SMC) and Sncg cells are depleted in the snRNA-seq 10x v3 B dataset, while Sncg cells are depleted in the MOP ST dataset. Besides, L6 IT Car3 and L4/5 IT are not provided in the snRNA-seq 10x v3 B dataset, thus these cell types are not considered in direct comparisons.

**Fig. 2.**
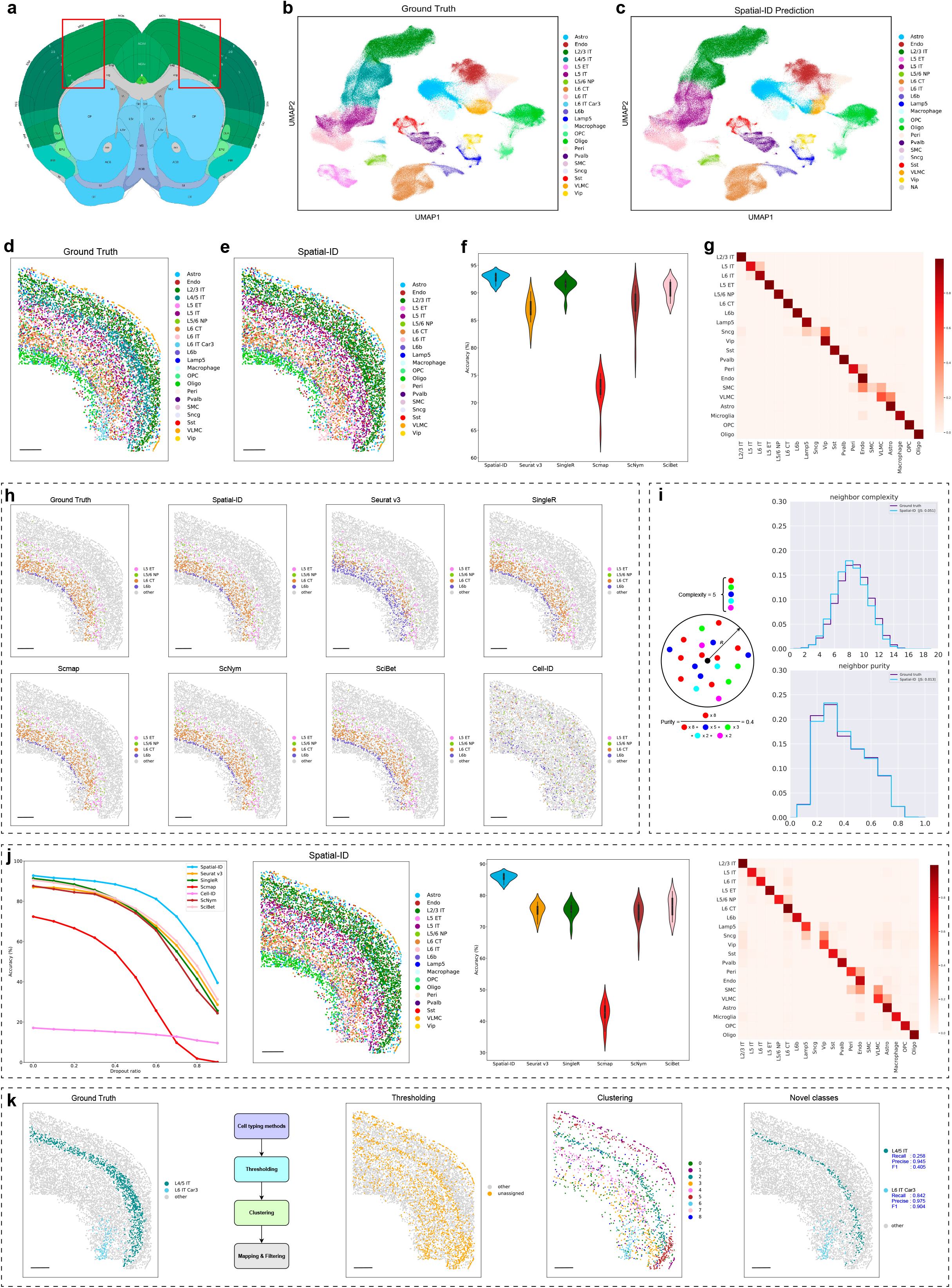
Application to mouse primary motor cortex dataset measured by MERFISH. (**a**) The MOP region annotations in the Allen CCF v3 (http://atlas.brain-map.org/). (**b**) Visualization of the ground truth cell types using UMAP embedding. (**c**) Visualization of the Spatial-ID predictions using UMAP embedding. (**d**) Spatial organization of the ground truth cell types in a coronal slices (slice153). Bar scale 400um. (**e**) Spatial organization of the Spatial-ID prediction in **d**. Bar scale 400um. (**f**) The comparison of mean accuracy. Notably, the mean accuracy of Cell-ID is far below those shown and is therefore not shown. (**g**) The confusion matrix shows the fraction of cells from any ground truth cell type predicted by Spatial-ID. The vertical axis lists the ground truth cell types, and the horizonal axis lists the cell types predicted by Spatial-ID. (**h**) Spatial organization of the ground truth of L5 ET, L5/6 NP, L6 CT and L6b neurons, and the prediction of Spatial-ID and control methods. Spatial-ID shows the highest accuracy and better spatial aggregation of each predicted cell type than the control methods. Bar scale 400um. (**i**) Spatial-ID shows high consistent of neighborhood complexity and neighborhood purity with ground truth. The neighborhood complexity of a given cell is defined as the number of different cell types presented within a neighborhood of 100 μm in radius. The neighborhood purity of a given cell is defined as the fraction of the most abundant cell type to all cells in the neighborhood of 100 μm in radius. (**j**) Simulations of different gene dropout rates. From left to right, the comparison of mean accuracy at different gene dropout rates, spatial organization of the Spatial-ID prediction at the dropout rate of 0.5, the comparison of mean accuracy at the dropout rate of 0.5, the confusion matrix of Spatial-ID prediction at the dropout rate of 0.5 are shown in sequence. Spatial-ID shows more robustness for gene dropout, because the degradation of Spatial-ID is less than that of the top control methods. Bar scale 400um. (**k**) New cell type discovery. From left to right, spatial organization of the ground truth of L4/5 IT and L6 IT Car3 neurons, a pipeline of new cell type discovery, spatial organization of unassigned cells derived from thresholding, spatial organization of clusters derived from clustering for unassigned cells, and the finally found novel cell types (i.e., L4/5 IT and L6 IT Car3) are shown in sequence. Bar scale 400um.

Compared with the control methods, Spatial-ID could effectively identify the cell types (Fig. 2c) and achieve better performance (Fig. 2f). On all 12 MERFISH SRT samples, Spatial-ID achieves the highest mean accuracy 92.75% (Fig. 2f), and the differences with the control methods are very significant (Wilcoxon test p-value<<0.001 for all other methods). Besides, Spatial-ID achieves the highest mean weighted F1 score 0.9209 (Extend Fig. 2a), where weighted F1 score of each sample is calculated by weighted averaging the F1 score of each cell type, in order to mitigate the effects of cell type imbalance. In addition, Cell-ID achieves mean accuracy 17.08% and mean weighted F1 score 0.1521 that is far below other methods and is therefore not shown in Fig. 2f and Extend Fig. 2a. From this, each cell in MOP SRT dataset have both a predicted cell type from the snRNA-seq 10x v3 B dataset and a MERFISH ground truth label. Cells in MOP SRT dataset are grouped based on their MERFISH ground truth labels, and then the fraction of predicted cell type for each group is determined (Fig. 2g and Extend Fig. 2d). Except the smooth muscle cells (SMC) and Sncg cells, Spatial-ID achieves outstanding recall rates on other cell types. The depleted SMC and Sncg cell types in the snRNA-seq 10x v3 B dataset lead to weaker identification ability of models on these cell types, because there are no sufficient samples to train models. Moreover, the integration of spatial information enables Spatial-ID to reveal that neuronal cells have specific spatial organization pattern (Fig. 2e). For the excitatory neurons, we can observe a laminar appearance for the overall cellular organization along the direction of the cortical depth, especially the IT neurons (Extend Fig. 1b). Obviously, the L2/3 IT, L5 IT, L6 IT neurons, identified by Spatial-ID, appear as discretely laminar cell populations, and L5 ET, L5/6 NP, L6 CT and L6b neurons also appear as discrete cell populations (Fig. 2h). However, IT neurons and other types of excitatory neurons partially overlap in space, and more overlapping with inhibitory neurons and non-neurons lead to a high level of local cellular heterogeneity. To describe the spatial intermixing of different cell populations, we count the neighborhood cells of each cell to calculate the neighborhood complexity^44^ and the neighborhood purity^44^, and then estimate the probability distribution for each identified cell type. According to the ground truth, the distribution of neighborhood purity presents very similar characteristic (Jensen-Shannon distance: 0.013), and the distribution of neighborhood complexity presents a slight shift to the lower complexity (Jensen-Shannon distance: 0.052) (Fig. 2i). Compared with the control methods (Extend Fig. 2b, Extend Fig. 2c, Extend Fig. 3 and Extend Fig. 4), Spatial-ID also achieves high consistent of neighborhood complexity and neighborhood purity with ground truth.

Different SRT technologies usually have different rates of gene capture, especially the spot-based SRT technologies. To verify the robustness of Spatial-ID in the application of SRT datasets with different gene dropout rates, we conduct simulation experiments on MOP SRT dataset by randomly discarding part of genes. Under the same configuration, Spatial-ID could achieve better performance of cell type identification than the control methods (Fig. 2j). Especially in the low dropout rates (e.g., less than 0.6), the performance degradation of Spatial-ID is less than that of the top control methods (Fig. 2j and Extend Fig. 5a). Specifically, on all 12 MERFISH ST samples, Spatial-ID achieves the highest mean accuracy 85.76% at the dropout rate of 0.5 (Fig. 2j), and the differences with the control methods are very significant (Wilcoxon test p-value<<0.001 for all other methods). While Spatial-ID achieves the highest mean weighted F1 score 0.8466 at the dropout rate of 0.5 (Extend Fig. 5b). These results suggest that the spatial information is not only beneficial to help cell type identification, but also improve the robustness of Spatial-ID against the variation of gene dropout (Fig. 2j, Extend Fig. 5c and Extend Fig. 5d). Thus, Spatial-ID shows a promising perspective to transfer knowledge from available reference datasets (e.g., scRNA-seq datasets or other SRT datasets), even if their gene dropout rates differ from that of target datasets.

L4/5 IT and L6 IT Car3 neurons are not provided in the snRNA-seq 10x v3 B dataset, thus Spatial-ID predicts the L6 IT Car3 neurons as L6 IT neurons, and L4/5 IT neurons as L2/3 IT and L5 IT neurons (Fig. 2c and Fig. 2e). The ground truth of MOP SRT dataset shows that L4/5 IT neurons populate in the continuous region between L2/3 IT and L5 IT neurons (Fig. 2d), and L6 IT Car3 neurons populate near the L6 IT neurons at L6 layer (Fig. 2d). These results suggest that the gene expression profiles of L4/5 IT neurons present gradual transition between the L2/3 IT and L5 IT neurons, and the characteristics of L6 IT Car3 neurons are similar to the L6 IT neurons (Fig. 2c). To further distinguish these new cell types, we conduct an experiment that some cell types are presented in the MOP SRT dataset but unseen in the snRNA-seq 10x v3 B dataset. Based on the prediction results of Spatial-ID, a pipeline of new cell type discovery (Fig. 2k) is performed to determine whether there are new cell types in the MOP SRT dataset, including thresholding, clustering, mapping and filtering. Finally, the discovery of L4/5 IT and L6 IT Car3 neurons achieves F1 score of 0.405 and 0.904, respectively (Fig. 2k).

### Application to mouse hypothalamic preoptic region dataset measured by MERFISH

For 3D SRT dataset, we also perform a quantitative comparison on the mouse hypothalamic preoptic region (1.8mm×1.8mm×0.6mm, Fig. 3a) dataset^44^ measured by MERFISH. This dataset measures a panel of 155 genes. We select 3 samples with naive behavior, including total 213,192 cells collected from 2 female mice and 1 male mouse (Fig. 3b). Each sample (Bregma 0.26 to −0.29) contains 12 slices with 50um interval. The reference scRNA-seq dataset (GSE113576)^44^ is collected from the hypothalamic preoptic region (∼2.5mm×2.5mm×1.1mm) across 3 replicates of an adult female mouse and a male mouse, including 31,299 cells and 27,998 genes. The delineation of major cell classes includes inhibitory neurons, excitatory neurons, microglia, astrocytes, immature oligodendrocytes, mature oligodendrocytes, ependymal cells, endothelial cells, macrophages and mural cells.

**Fig. 3.**
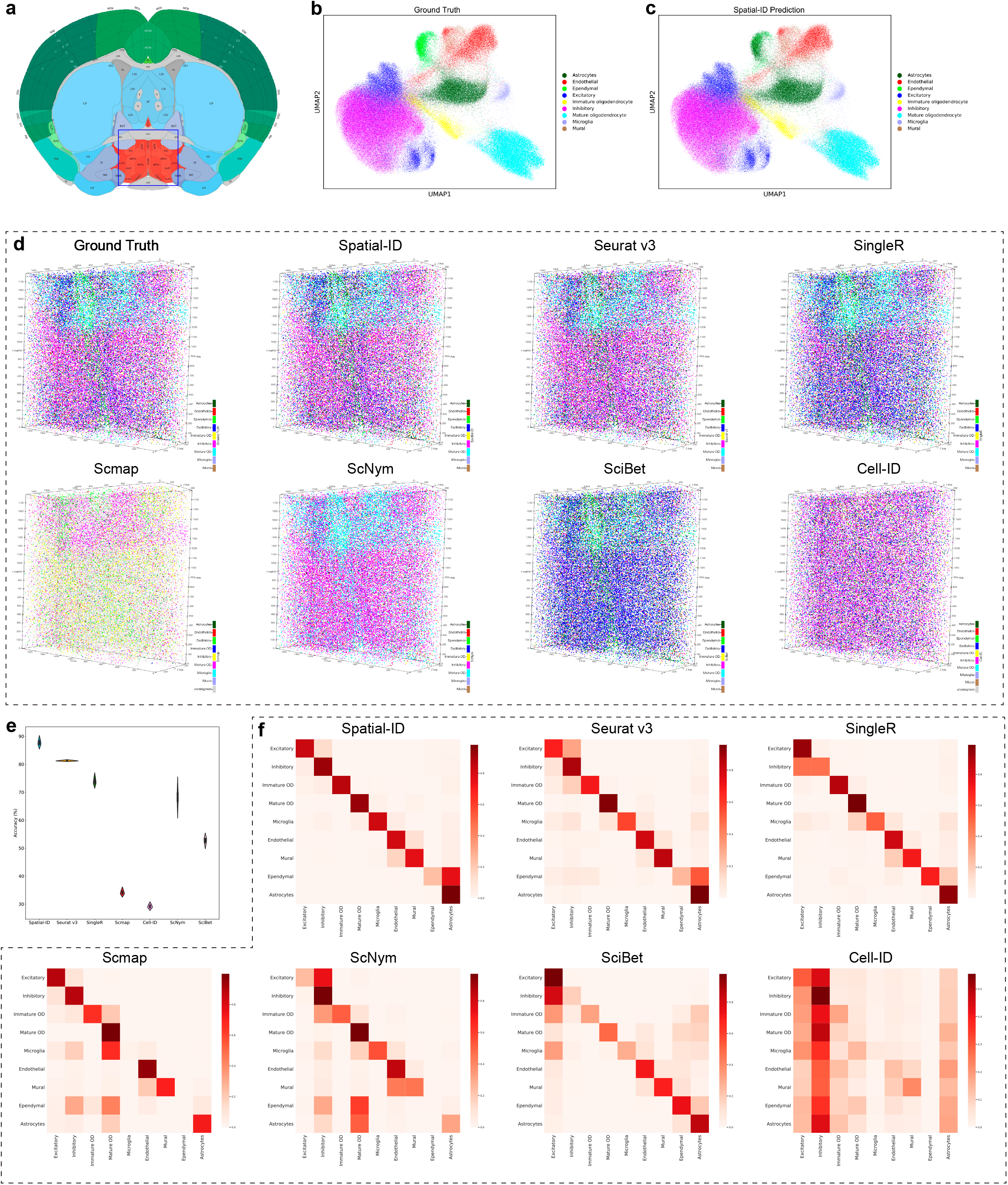
Application to mouse hypothalamic preoptic region dataset measured by MERFISH. (**a**) The mouse hypothalamic preoptic region annotations in the Allen CCF v3 (http://atlas.brain-map.org/). (**b**) Visualization of the ground truth cell types using UMAP embedding. (**c**) Visualization of the Spatial-ID predictions using UMAP embedding. (**d**) 3D spatial organization of the ground truth cell types of a sample with naive behavior, and the predictions of Spatial-ID and control methods. Scale unit (um). (**e**) The comparison of mean accuracy. (**f**) The confusion matrixes show the fraction of cells from any ground truth cell type predicted by Spatial-ID and control methods. The vertical axis lists the ground truth cell types, and the horizonal axis lists the predicted cell types.

Spatial-ID achieves the highest mean accuracy 87.74% (Fig. 3c and Fig. 3e) that significantly outperforms the control methods (Wilcoxon test p-value<<0.001 for all other methods) on all 3 samples with naive behavior (Fig. 3d and Fig. 3e). Besides, Spatial-ID achieves the highest mean weighted F1 score 0.8773 (Extend Fig. 7a). These results suggest that Spatial-ID can be effectively applied on 3D SRT dataset. The cell types annotated by Spatial-ID (Fig. 3f) show better correspondence to ground truth than those annotated by control methods. Specifically, in the 2D views (Extend Fig. 6a and Extend Fig. 6b), the annotation results of Spatial-ID show more obvious advantage than other control methods.

### Application to mouse spermatogenesis dataset measured by Slide-seq

Next, we perform a quantitative comparison on the mouse spermatogenesis dataset^45^ (Fig. 4a) measured by Slide-seq. The mouse spermatogenesis SRT dataset is acquired from three leptin-deficient diabetic (ob/ob) mice and three wild-type (WT) mice, including 207,335 cells in total and 24,105 genes in common. All cells are divided into 9 testicular cell types^69^: elongating/elongated spermatid (ES), round spermatid(RS), spermatocyte(SPC), spermatogonium(SPG), Endothelial, Sertoli, Leydig, Myoid and Macrophages (Fig. 4a). Testicular cell types are organized in a spatially segregated fashion at the level of seminiferous tubules. By comparison in previous study^45^, diabetic induces testicular injuries through disrupting the spatial structures of seminiferous tubules and changing the expression pattern of many genes at molecular level (Fig. 4e). The reference scRNA-seq dataset (GSE112393)^48^ includes 34,633 cells and 37,241 genes from the adult mouse testis.

**Fig. 4.**
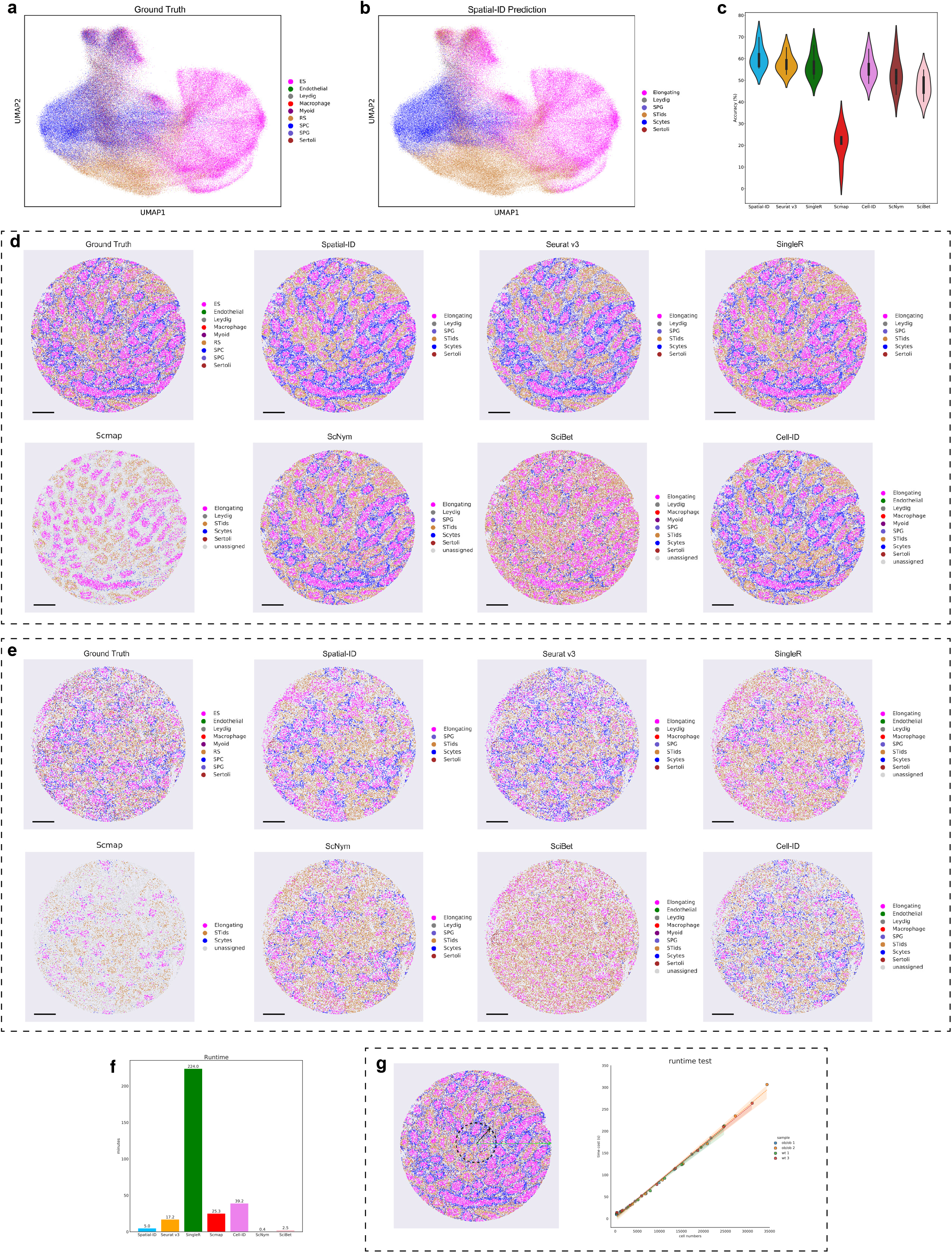
Application to mouse spermatogenesis dataset measured by Slide-seq. **(a)** Visualization of the ground truth cell types using UMAP embedding. (**b**) Visualization of the Spatial-ID predictions using UMAP embedding. (**c**) The comparison of mean accuracy. (**d**) Spatial organization of the ground truth cell types of a wild-type sample, and the predictions of Spatial-ID and control methods. Bar scale 400 um. (**e**) Spatial organization of the ground truth cell types of an ob/ob sample, and the predictions of Spatial-ID and control methods. Bar scale 400 um. (**f**) The average time cost per sample of Spatial-ID and control methods in this mouse spermatogenesis dataset. The comprehensive results for all SRT datasets in this studied can be found at Table 3. (**g**) The running efficiency analysis. The left one shows the scheme of field view sampling. The right one shows that the runtime of Spatial-ID increases linearly as the number of cells increases.

**Table 3.** The average time cost (mins) per sample of Spatial-ID and control methods. Notably, the supervision-based methods (i.e., ScNym and SciBet) do not count the time of models training. For the self-collected mouse brain hemisphere, the average time cost per sample of the control methods are compared in the configuration of 747 marker genes, because correlation-based methods require huge computational resources for tens of thousands of genes.

| SRT datasets | average cells per sample, genes | Spatial-ID | Seruat v3 | SingleR | Scmap | Cell-ID | ScNym | SciBet | Used Cells in Reference datasets |
| --- | --- | --- | --- | --- | --- | --- | --- | --- | --- |
| Primary motor cortex (MOP) | 22,746 cells, 254 genes | 2.4 | 51.7 | 50.3 | 12.5 | 75.9 | 0.2 | 0.3 | 159,738 cells |
| Hypothalamic preoptic region (3D) | 71,064 cells, 155 genes | 9.1 | 16.3 | 51.3 | 19.0 | 32.7 | 0.1 | 0.2 | 31,299 cells |
| Mouse spermatogenesis | 34,556 cells, 24015 genes | 5.0 | 17.2 | 224.0 | 25.3 | 39.2 | 0.4 | 2.5 | 34,633 cells |
| Mouse brain hemisphere (3D) | 59,025 cells, 747 genes | 6.1 | 31.8 | 285.7 | 66.0 | 84.4 | 0.1 | 0.2 | 113,488 cells |

The quantitative comparison on mouse spermatogenesis dataset also demonstrates the superiority of Spatial-ID for cell type identification molecularly (Fig. 4c, Fig. 4d and Fig. 4e). On all 6 SRT samples, Spatial-ID achieves the highest mean accuracy 60.45% (Fig. 4c), and the differences with the control methods are very significant (Wilcoxon test p-value<<0.001 for all other methods). These results suggest that Spatial-ID can also effectively handle the spot-based Slide-seq dataset with tens of thousands of genes, even if the area of a spot span more than one cell. Besides, Spatial-ID achieves a mean weighted F1 score 0.55 (Extend Fig. 7b). In this comparison, all 24,015 common genes are used in Spatial-ID and control methods. Interestingly, Cell-ID presents outstanding performance in this SRT dataset, which achieves the highest mean weighted F1 score 0.5727 (Extend Fig. 7b) that indicates Cell-ID requires abundant genes for cell type annotation.

Moreover, we compare the runtime of Spatial-ID and control methods on this SRT dataset (Fig. 4f). The running efficiency of supervision-based methods (i.e., Spatial-ID, ScNym and SciBet) is much higher than that of correlation-based methods. The same results can be obtained for other SRT datasets (See Table 3). To further analyze the running efficiency of Spatial-ID on different number of cells, we crop 10 percent to 100 percent radius field views of 4 samples (other 2 samples have non-circular field views) in this SRT dataset (Fig. 4g), respectively. Then, we statistically analyze the running efficiency of Spatial-ID on these cropped field views. As the number of cells increases, the runtime of Spatial-ID increases linearly (Fig. 4g).

### Application to large field mouse brain hemisphere dataset measured by Stereo-seq

Many currently available sequencing-based SRT technologies such as Slide-seq, DBiT-seq^49^ and HDST do not have single-cell spatial resolution, where each spot contains 1 to 10 cells. With the continued improvement of spatial resolution, newly emerging SRT technologies such as Seq-Scope^50^ and Stereo-seq can produce high-throughput subcellular SRT data with a large number of cells in large field tissues in subcellular spatial resolution. Here, we generate single-cell spatial gene expression profiles of 3 adjacent coronal sections (10μm thick, without intervals) along the anterior-posterior axis (Bregma −3.56 to −3.66) of right mouse brain hemisphere (Fig. 5a) using Stereo-seq. After the standard data processing and quality control (See Methods), 140,816 cells are retained for these 3 sections. The single-cell mouse brain atlas of cell types from the Linnarsson Lab^51^ is employed as the reference dataset. After selecting the cell types of reference dataset that located in brain sections of our samples, a subset of 113,488 cells belonging to 152 cell types with a total of 747 marker genes is used as training set of the proposed Spatial-ID.

**Fig. 5.**
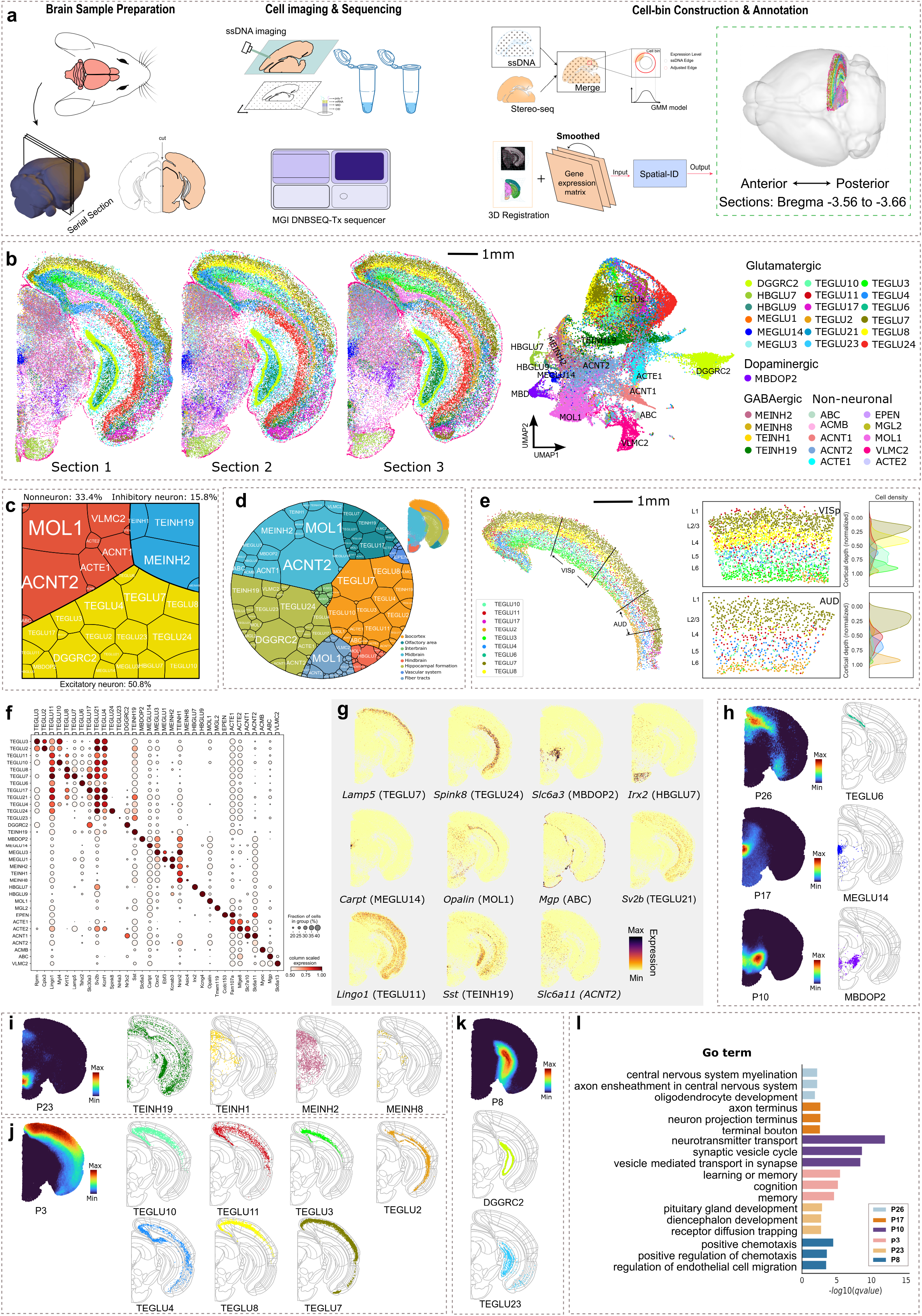
Application to large field mouse brain hemisphere dataset measured by Stereo-seq. **(a)** The workflow of data acquisition, data processing and cell type annotation. **(b)** Cell type annotation of Spatial-ID for the 3 adjacent sections (Bregma −3.56 to −3.66 mm), and UMAP visualization. **(c)** A Voronoi treemap shows the composition of excitatory neurons, inhibitory neurons and non-neuronal cells among the 3 sections. Every tile denotes one cell type and its size represents cell number. **(d)** A Voronoi diagram shows cell type organization among distinct brain regions of the 3 sections. Every tile is colored by its populated ABA functional region and its size represents cell number. **(e)** Spatial organization of the cortical pyramidal neurons, i.e., TEGLU2, TEGLU3, TEGLU4, TEGLU6, TEGLU7, TEGLU8, TEGLU10, TEGL11 and TEGLU17 in the Section 3. Cells in the VISp and AUD region are individually presented in the middle panel. The right panel shows the kernel density estimate plots for the corresponding cell types along the normalized cortical depth. **(f)** The expression dot plots show the gene expression specificity of typical marker genes for identified cell types. Dot size represents the proportion of expressing cells and color indicates average expression level in each identified cell type. **(g)** Spatial distributions of selected marker genes show the number of transcripts captured by Stereo-seq. **(h)** The spatial gene patterns consist of type-specific genes (Section 3, visualized with pattern scores). The right panel shows the corresponding identified cell types together with the ABA spatial anatomical functional regions. **(i-k)** The spatial gene patterns consist of region-specific genes from diverse identified cell types (Section 3). The corresponding identified cell types are illustrated on the right. **(l)** Top three highly enriched GO terms for each spatial gene pattern in **h-k**.

Based on the predictions of Spatial-ID, the identified cell types of 3 coronal sections present a high consistency (Fig. 5b) that an average of 99% cells in each section are assigned to their common cell types (Extend Fig. 8e). Besides, by color-coding our identified cell types in low dimension feature space (i.e., UMAP), most identified cell types congregate into separable communities (Fig. 5b). For example, DGGRC2, VLMC2, and MBDOP2 can be easily segregated because they are driven by large differences of gene expression. Some similar cell types populate in mixed communities, such as the cortical pyramidal neuron (TEGLUs). According to the cell type taxonomy of reference dataset, 65,174 cells (50.8%) are identified as excitatory neurons, 20,267 cells (15.8%) are identified as inhibitory neurons, and 42,840 cells (33.4%) are identified as non-neuronal cell types (Fig. 5c). Specifically, most of the identified excitatory neurons are telencephalon projecting neurons with glutamatergic neurotransmitter (TEGLU) and populate in cerebral cortex and hippocampus, such as TEGLU4, TEGLU7 and TEGLU8 in cerebral cortex (Fig. 5e and Fig. 5j), TEGLU24 (Extend Fig. 9a) and DGGRC2 (Fig. 5k) in hippocampus. The other excitatory neurons with glutamatergic neurotransmitter (MEGLU, HBGLU) populate in midbrain and hindbrain. The identified inhibitory neurons mainly consist of TEINH19 and MEINH8 (Fig. 5i), where the identified TEINH19 neurons scatter across cortical layers and hippocampus CA3 region and the identified MEINH8 populate in midbrain. The identified non-neuronal cells exhibit a dispersed distribution throughout the mouse brain hemisphere, such as ACNT2 (non-telencephalon astrocyte cells) in midbrain, hindbrain and fiber tracts (Extend Fig. 9a), VLMC2 (vascular leptomeningeal cells) at the interface of the brain structure (Extend Fig. 9a).

As the visualization of cell type annotation in Fig. 5b, most of identified cells show a separable spatial organization. To further reveal the anatomical functions of identified cell types throughout distinct brain regions, the entire right mouse brain hemisphere can be roughly split into several spatial anatomical functional regions according to the Allen Brain Atlas^25^ (ABA; https://atlas.brain-map.org/), including isocortex, hippocampal formation, olfactory area, midbrain, hindbrain, interbrain, fiber tracts and vascular system. By quantifying the cells among these 8 regions, we can observe that different functional regions have different combinations of the identified cell types (Fig. 5d). For example, in the isocortex region (Fig. 5e), the identified cortical pyramidal neurons (TEGLU7, TEGLU8, TEGLU10, TEGLU4, TEGLU3, TEGLU2, etc.) display a layered laminar appearance along the direction of the cortical depth^52–54^. Moreover, we further illustrate a continuous gradient of cells along the cortical depth from L2/3 to L6 in the VISp and AUD regions (Fig. 5e). The cell type compositions of the VISp and AUD regions have significant differences, where fewer TEGLU3 and TEGLU8 populate in the AUD region than the VISp region (Fig. 5e).

For the mainly identified cell types, we further analyze the gene expression specificity of typical marker genes provided by the reference dataset^51^ (Fig. 5f). Most of these marker genes have the highest expression in their corresponding cell types that have a relatively high fraction, e.g., *Lamp5* of TEGLU7, *Spink8* of TEGLU24, Slc6a3 of MBDOP2, *Irx2* of HBGLU7, *Carpt* of MEGLU14, *Opalin* of MOL1, *Mgp* of ABC (Fig. 5g), etc. Moreover, several marker genes, such as *Sv2b* of TEGLU21, *Lingo1* of TEGLU11 and *Sst* of TEINH19, present continuous expressions across the identified neurons in cerebral cortex and hippocampus (Fig. 5g). Interestingly, we observe the ACNT2 maker gene *Slc6a11* expressed higher in ACNT1, another subclass of non-telencephalon astrocytes, than in ACNT2 (Fig. 5g). These observations may be derived from the continuous variation among neighborhood subclasses or the combinatorial expression of marker genes.

We further investigate the spatially varying genes indicative of the underling cell types, and their grouped spatial patterns. A total of 30 specific spatial gene patterns are detected by Hotspot (Extend Fig. 9b) and 6 of them are illustrated in Fig. 5h, Fig. 5i, Fig. 5j and Fig. 5k. Notably, a spatial gene pattern may consist of type-specific genes from an individual identified cell type (Fig. 5h), but it may also be constituted by region-specific genes from diverse identified cell types (Fig. 5i, Fig. 5j and Fig. 5k). Specifically, the spatial gene pattern P26 is detected in the retrosplenial area of layer 2 (Fig. 5h), which includes *Tshz2* (Extend Fig. 9d), one of the marker genes of the cortical projection neurons TEGLU6 that are identified in this region (Fig. 5h). The GO-based enrichment result indicates that the spatial gene pattern P26 may involve in myelination and axon ensheathment of central nervous system, possibly supporting a role for retrosplenial cortex in spatial coding, memory formation, and information integration^71,72^ (Fig. 5l). In the midbrain, the obviously spatial gene pattern P17 (Fig. 5h), contains gene *Ucn*, *Slc5a7*, *Chodl*, etc (Extend Fig. 9d), significantly enriches in the sub-region of dorsal raphe nucleus (DRN). Accordingly, the identified MEGLU14 neurons (marked genes: *Cartpt, Ucn and Chodl*) specifically populate in this region (Fig. 5h). DRN has been implicated in disorder of anxiety, reward processing, as well as social isolation^73,74^. Here, we find that these DRN-specific genes are highly enriched at axon terminus, neuron projection terminus and terminal bouton (Fig. 5l), which are specialized to release neurotransmitters to transmit impulses between neurons. Another spatial gene pattern P10 is detected in the sub-regions of ventral tegmental area and subtantia nigra (SNr, Fig. 5h), contains genes *Slc6a3*, *Slc18a2* and *Th*, etc (Extend Fig. 9d). This spatial gene pattern corresponds to identified MBDOP2 neurons (marked genes: *Slc6a3* and *Chrna6*), which are dopaminergic neurons in midbrain that have been reported to be associated with the genetic risk of neuropsychiatric disorders^59^, for example Parkinson’s disease. The further GO-based enrichment result indicates that these enriched genes may involve in the regulation of neurotransmitter levels (Fig. 5l), revealing the relationship between gene expression of MBDOP2 neurons and neuropsychiatric disorders again. Besides, serval identified spatial gene patterns do not indicate to specific cell types, such as P23 (Fig. 5i), P3 (Fig. 5j) and P8 (Fig. 5k). Particularly, P23, which is highly concentrated in the interpeduncular nucleus (IPN) of ventral midbrain, consists of several GABAergic neuron-associated genes (e.g., *Otp*, *Pax7*, *Gad1*, *Gad2*, *Slc32a*), suggesting its representative role for inhibitory neurons. As expected, we can observe that the identified inhibitory neurons including TEINH19, MEINH2, MEINH8 and TEINH1 populate in this area (Fig. 5i). Previous studies found that IPN was the critical brain area associated with the reinforcing effects of nicotine^62^. The GO-based enrichment result reveals these spatially enriched genes are highly related with pituitary gland and diencephalon development and receptor diffusion trapping (Fig. 5l). P3 is highly concentrated in the isocortex (Fig. 5j), which indicates the identified cortical pyramidal neurons (TEGLUs shown in Fig. 5e), and significantly enriched in GO terms of learning, memory and cognition (Fig. 5l). P8 is found to be involved in positive chemotaxis, and mainly constituted by genes from hippocampal neurons DGGRC2 and TEGLU23 (Fig. 5k).

## Discussion

In this work, we propose Spatial-ID to perform cell type annotation for SRT datasets molecularly. We first conduct a series of comparisons on 3 public available SRT datasets with different characteristics (See Table 1). By comparing the accuracy and weighted F1 score calculated from predictions and ground truth, the proposed Spatial-ID demonstrates superior performance than the state-of-the-art methods (See Table 2) on two FISH-based SRT datasets with hundreds of genes measured by MERFISH technology and a spot-based SRT dataset with tens of thousands of genes measured by Slide-seq technology. Specifically, the better performance of Spatial-ID on the 3D FISH-based SRT dataset (i.e., mouse hypothalamic preoptic region) further confirms that Spatial-ID can be effectively applied to 3D SRT dataset. Moreover, the comparisons on the randomly gene dropout simulations of the FISH-based SRT dataset (i.e., mouse primary motor cortex) additionally demonstrates the better robustness of Spatial-ID against the variation of gene dropout. Besides, the extended experiment of new cell type discovery shows the adaptability of Spatial-ID in the situation of transfer learning from an incomplete reference dataset. These results suggest that embedding spatial information can not only improve the accuracy of cell type identification substantially, but also improve the robustness against the variation of sequencing technologies, such as different gene dropout rates, different number of measured genes. In addition, the running efficiency of Spatial-ID on all SRT datasets (See Table 3) is much higher than that of correlation-based methods (i.e., Seurat v3, SingleR, Scmap and Cell-ID).

As the application of Spatial-ID on the large field mouse brain hemisphere dataset measured by Stereo-seq technology, we investigate the spatial organization of identified cell types that show high consistency with previous studies at spatial anatomical structure level, such as Allen Brain Atlas. By analyzing the gene expression specificity of typical marker genes reported in reference dataset, i.e., the single-cell mouse brain atlas of cell types from the Linnarsson Lab^51^, most of these pre-defined marker genes have the highest expression in their corresponding cell types. Besides, by mapping the identified cell types with identified spatial gene patterns, the significant GO terms of the spatial gene patterns further reveal the functions and underlying biological processes of the identified cell types in mouse brain. Therefore, Spatial-ID shows a promising perspective to build a large-field spatial transcriptomic brain atlas.

In principle, the proposed Spatial-ID is a reference-based supervised cell typing method, therefore Spatial-ID is influenced by the characteristics of the reference datasets such as cell number, cell heterogeneity, the gene set used as features. For example, the depleted cell types in reference dataset lead to insufficient knowledge transfer of Spatial-ID, that may present weaker identification ability on these cell types. Besides, the Spatial-ID is influenced by the hyperparameters of constructing graph network such as the number of nearest cells and coefficient of inverse distance weight, that also indicates the intensity of spatial information used. The excessive use of spatial information may lead to reduce differences between local cells. Therefore, we anticipate that the more spatial information used enables the Spatial-ID to enhance the identification of locally enriched cell types and inhibit the identification of rare cell type. In addition, the Spatial-ID demonstrates well robustness against the variation of gene dropout, but it is inevitably influenced by the high dimensionality and biological noise associated with SRT sequencing technologies. Beside optimizing the data preprocessing, integrating the SRT datasets with scRNA-seq datasets, that have been determined with rich mRNA transcripts, or other modalities, such as microscopy data and proteomics data, is potential to mitigate the sequencing noise.

## Methods

### Overview of Datasets

Our proposed Spatial-ID was performed on different SRT datasets generated by MERFISH, Slide-seq, Stereo-seq SRT technologies. For each group of applications, we also collected reference scRNA-seq datasets. Except the self-collected large field mouse brain hemisphere dataset measured by Stereo-seq, all other SRT datasets and reference scRNA-seq datasets are publicly available, as shown in Table 1. Notably, the mouse hypothalamic preoptic region dataset measured by MERFISH and the self-collected mouse brain hemisphere measured by Stereo-seq are 3D SRT datasets. In addition, the field size of the self-collected mouse brain hemisphere is much larger than other public datasets.

### Data collection of large field mouse brain hemisphere

The large field mouse brain hemisphere dataset was measured by Stereo-seq^17^ (Fig. 5a) Briefly, three adjacent coronal sections (10μm thick, without intervals) along the anterior-posterior axis (Bregma −3.56 to −3.66) were cut from the brain of a 5-week-old C57BL/6J male mouse (Fig. 5a). First, these sections were adhered to the DNA nanoball (DNB, i.e., spot) patterned Stereo-seq chip surface, and then were stained and scanned into ssDNA images for cellular localization. Each DNB on Stereo-seq chip contains a 25 bp randomly barcoded sequence as coordinate identity (CID) for its unique spatial location, a 10 bp molecular identifier (MID) and a polyT for *in situ* RNA capture, having a size of 220 nm in diameter and 500 nm center-to-center distance. Next, tissue permeabilization, in situ reverse transcription, amplification, library construction and sequencing were performed according to the protocol of Stereo-seq technology.

### Data processing of large field mouse brain hemisphere

CID sequences were first parsed to the spatial coordinates of the DNB patterned silicon chip, allowing 1 base mismatch to correct for sequencing and PCR errors. Qualified reads (Q score ≥ 10) were then aligned to the mouse reference genome Ensembl GRCm38 v86 via STAR. Mapped reads with MAPQ ≥ 10 were annotated, and counted by handleBam (available at https://github.com/BGIResearch/handleBam). A resulting CID-containing expression profile matrix was thus generated for each section. To assign the captured RNAs to individual cells, we transformed the CID-containing gene expression matrix to an image by summing the MID counts at every pixel, and aligned with the corresponding ssDNA image based on patterned track lines in both images. Cell segmentation was performed on the registered ssDNA images by a UNet-like deep convolutional neural network (Extend Fig. 8a). To recall the genes from the cytoplasm surrounding the nucleus, a Gaussian mixture model was employed to adjust the cell boundaries. Thus, a single cell-based gene expression matrix was generated for each section by aggregating the spots included in each cell (Extend Fig. 8a). To facilitate the analysis of anatomical function of mouse brain, these 3 sections were aligned to the three-dimensional (3D) standard Allen Brain Atlas (Extend Fig. 8b) by Wholebrain (https://github.com/tractatus/wholebrain).

### QC and Gaussian smoothing for large field mouse brain hemisphere

The 3 adjacent coronal sections captured 53,310 cells, 61,910 cells and 61,857 cells, respectively. Pearson coefficient was used to evaluate the consistency between them (Extend Fig. 8c). Low-quality cells and genes were discarded according to the following quality control (QC) criteria: 1) cells with total counts lower than 300 and higher than 98% quantile, 2) cells with percentage of mitochondrial genes larger than 10%, 3) genes presented in less than 10 cells. Thus, 41,766 cells, 48,721 cells, 50,329 cells were remained for the 3 adjacent coronal sections (∼434 genes per cell), respectively. Next, a Gaussian smoothing strategy was introduced to alleviate the noise and gene dropout associated with the Stereo-seq technology (Extend Fig. 8d). Specifically, the principal component analysis (PCA) was firstly applied cell-wise to reduce the raw gene expression matrix into a low dimension feature space. Then a number of nearest neighboring cells of each cell in the feature space were acquired, which were used to update the gene expression of current cell by Gaussian smoothing.

### Deep neural networks for transfer learning

The transfer learning strategy of Spatial-ID employs reference scRNA-seq datasets to train the deep neural network (DNN) model (Fig. 1a) that is used to generate the probability distribution *D* of each cell in SRT data. It should be noted that the DNN model only take the gene expression profile as input, whereas the available spatial information of SRT data is not used. The DNN provides nonlinear dimensionality reduction for the input gene expression matrix, that consists of 4 stacked fully connected layers, as well as a GELU layer (nonlinear activation function) and a dropout layer followed by each fully connected layer in sequence. Moreover, to alleviate the class imbalance of cell types, the loss function of Focal Loss is employed in DNN models training.

To perform transfer learning, the gene sets of reference scRNA-seq dataset and SRT dataset should be matched. We first compare the measured genes to find the common gene set. If the reference scRNA-seq datasets or SRT datasets provide marker genes of cell types, we further select the subset of marker genes from the common gene set. This process can simplify the implementation of transfer learning. Then, the raw counts of selected genes from reference scRNA-seq dataset and SRT dataset are normalized to unit length vector for each cell. If there is no marker gene available (e.g., mouse spermatogenesis), we additionally perform a stage of *log*1*p* operation to raw counts of genes before the normalization, that intends to inhibit the negative effects of very highly expressed genes.

### Spatial neighbor graph construction for spatial information

To perform spatial embedding, we construct a spatial neighbor graph to represent the spatial relationships between neighboring cells (Fig. 1b). The spatial neighbor graph consists of nodes and edges, where a node represents a cell and an edge represents the relationship of a pair of neighboring cells. To better characterize the relationships, we calculate the Euclidean distance between current cell and neighboring cells using the spatial coordinates. Because the behavior of an individual cell is mediated by the ligand-receptor interactions with its neighboring cells in local tissue microenvironment, so closer distance indicates more closer relationship. For each selected neighbor, we calculate the weight negatively associated with its Euclidean distance by

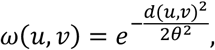

where *d*(*u*, *v*) denoted the Euclidean distance of a pair of neighboring cells, *0* denoted the decay coefficient. Specifically, we select the top *N* nearest neighbors (e.g., 30 in this study) of each cell to create the adjacency matrix, denoted by *A*, in which a cell *u* with top *N_u_* neighboring cells can be calculated by *A*(*u*, *v_i_*) = *w*(*u*, *v_i_*) *if v_i_ ∈ N_u_ else* 0.

### Deep autoencoder for latent representation learning

A deep autoencoder^39^ is used to learn encoded gene representation *X* through reducing the dimension of the gene expression matrix *I* (Fig. 1c). If the SRT datasets contains tens of thousands of genes, the gene expression matrix *I* is generated by extracting hundreds of principal components (e.g., 200) of principal component analysis (PCA). The encoder part consists of 2 stacked fully connected layers, as well as a batch normal layer, a ELU layer (nonlinear activation function) and a dropout layer followed by each fully connected layer in sequence. The decoder part consists of one fully connected layer and same followed layers as encoder. The deep autoencoder employs the mean squared error (MSE) loss function to maximize the similarity between the input gene expression matrix *I* and the output gene expression matrix *I’* reconstructed by decoder.

### Variational graph autoencoder for spatial embedding

A GCN^38^, i.e., variational graph autoencoder (VGAE)^40^, is used to embed spatial neighbor graph (Fig. 1d). Because the spatial neighbor graph contains large number of nodes for high-throughput cell-level SRT data, we employ sparse graph convolution layers in VGAE to accelerate the computation. The variational modification of the encoder-decoder architecture can introduce regularization in the latent space, thus improves the properties of spatial embeddings. The graph encoder takes encoded gene representations *X* from autoencoder and the adjacency matrix *A* as input, then generates the spatial embedding *s* as output. The graph encoder consists of 2 sparse graph convolution layers, as well as a RELU layer (nonlinear activation function) and a dropout layer followed by each graph convolution layer in sequence. The first sparse graph convolution layer is used to generate a lower-dimensional feature matrix. Next, the second sparse graph convolution layers generate a feature matrix *µ* and a feature matrix *logσ*^2^, respectively. Then the spatial embedding *S* is calculated using parameterization trick *S* = *µ* + *σ ∗ τ*, where *τ*∼*N*(0,1). The final latent representations *Z* are combined from the encoded gene representation *X* and the spatial embedding *S* by formula *Z* = *X* + *S*. Thereafter, the final latent representations *Z* are used to reconstruct the gene expression matrix *I’* in the autoencoder and the adjacency matrix *A’* in the VGAE. Specifically, the graph decoder adapts an inner product to reconstruct adjacency matrix *A’* from the spatial embedding *Z*. The loss function of the VGAE employs a cross-entropy loss to minimize the input adjacency matrix *A* and the reconstructed adjacency matrix *A’*, and a KL-divergence to measure the similarity between *q*(*Z*| *X*, *A*) and *p*(*Z*), where *p*(*Z*)∼*N*(0,1).

### Self-supervised learning using pseudo-labels

We additionally perform a self-supervised learning strategy to train a classifier using the final latent representations *Z* and pseudo-labels *L*. In this strategy, the DNN model is the teacher model and the classifier is the student model. Formally, the pseudo-labels *L* = {*l*_1_, …, *l_i_*, …, *l_n_*} are derived from the output *Y* = {*Y*_1_, …, *Y_i_*, … *Y_n_*} of last fully connected layer of the DNN model by a modified *softmax* layer

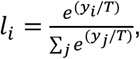

where *T* is the temperature parameter that adjusts the smoothness of distribution (Fig. 1b), and *L* = *D*, if *T* = 1. Obviously, a higher value for *T* produces softer distribution of pseudo-labels *L* over all classes. The softer distribution of pseudo-labels *L* could transfers more information from the reference scRNA-seq datasets.

### Simulation of different gene dropout rates

Many spot-based ST technologies usually have lower mRNA capture efficiency due to the limited number of probes. Therefore, the gene dropout rates of spot-based SRT datasets are usually higher than the state-of-the-art scRNA-seq technologies and FISH-based SRT technologies. To verify the adaptability of Spatial-ID in different gene dropout rates, we conduct simulation experiments on FISH-based mouse primary motor cortex SRT dataset (Fig. 2j). A random-gene-discard strategy, that randomly clears part of value in gene expression matrix, is employed to generate simulated SRT datasets. Then, we compare cell type annotation of Spatial-ID and control methods on these simulated SRT datasets.

### New cell type discovery

The discovery of new cell populations has biologically important implication for omics analysis. Technically, the new cell type discovery is an anomaly detection task, because the trainset does not contain the new cell types to be detected. To perform new cell type discovery, we introduce an extended postprocessing strategy for Spatial-ID that includes thresholding, clustering, mapping and filtering. For each cell, the maximum score *c_s_ ∈* [0,1] of predicted probability vector *C* = {*c*_1_, …, *c_i_*, …, *c_n_*_−1_}, 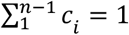 is first examined by a threshold. If the score *c_s_* of the cell is less than the threshold (e.g., 0.9 in Fig. 2k), the cell is recognized as an unassigned cell. After a set of unassigned cells are retrieved in the SRT dataset, a clustering stage is performed to distinguish these unassigned cells into different clusters in gene expression feature space, because the new cell type should populate in a relatively isolated area. Finally, through mapping analysis of the gene expression profiles between these clusters and known cell types, the matching clusters are recognized as known cell types and the mismatching clusters are recognized as new cell types.

### Identifying spatial gene patterns for large field mouse brain hemisphere

Hotspot^56^ is used to identify spatial gene patterns (Extend Fig. 9b). First, data binning is performed to reduce computational difficulty by dividing the x-y coordinates into grids covering an area of 50*50 DNB (bin50) and the transcripts of the same gene aggregated within each bin. Next, highly variable genes are found by toolkit scanpy^63^. Hotspot uses the number of 300 neighbors to create the spatial KNN graph, and then uses spatially-varying genes (FDR <= 0.05) to identify spatial gene patterns. In addition, clusterProfiler^57^ is used to perform GO enrichment analysis of the identified spatial gene patterns (Fig. 5j).

### Computational resources and runtime

All analyses presented in the paper are run in a workstation with 40 Gb RAM memory, 10 cores of 2.5 GHz Intel Xeon Platinum 8255C CPU, and a Nvidia Tesla T4 GPU with 8 Gb memory. The runtime of different cell typing methods in this workstation are shown in Table 3.

## Acknowledgements

This project has been deposited at the CNSA under the BioProject number CNP0002966.

## Author contributions

Conceptualization: Rongbo Shen, Lin Liu, Zihan Wu and Ying Zhang.

Project administration and supervision: Yuxiang Li, Xun Xu and Jianhua Yao.

Algorithm development and implementation: Zihan Wu and Rongbo Shen.

Public datasets collection, processing and application: Rongbo Shen, Zihan Wu and Zhiyuan Yuan.

Stereo-seq dataset collection, processing and application: Zihan Wu, Lin Liu, Ying Zhang, Rongbo Shen, Junfu Guo, Chao Zhang, Bichao Chen, Chao Liu, Jing Guo, Guozhen Fan, Yong Zhang.

Methods comparisons: Zihan Wu and Rongbo Shen.

Biological interpretation: Lin Liu, Rongbo Shen and Ying Zhang.

Project coordination: Fang Yang and Bichao Chen.

Manuscript writing and figure generation: Rongbo Shen and Lin Liu.

Manuscript reviewing: Zhiyuan Yuan, Fang Yang and Jianhua Yao.

All authors approved the manuscript.

## Competing interests

The authors declare no competing interests.

---

**Extend Fig. 1.**
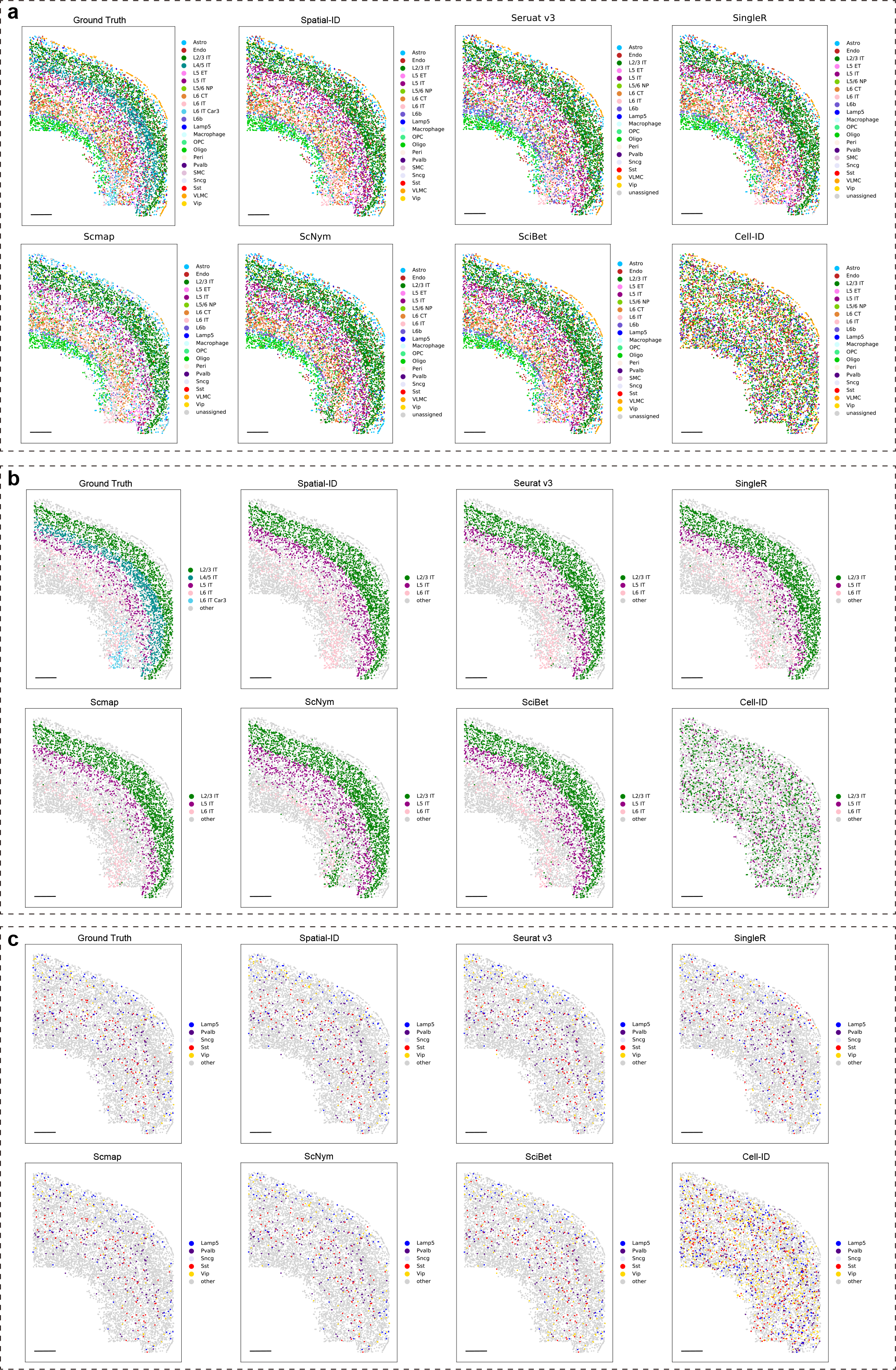
Spatial organization of ground truth and predictions of different cell typing methods. Bar scale 400um. (**a**) All cell types. (**b**) Intra-telencephalic neurons (L2/3 IT, L4/5 IT, L5 IT, L6 IT and L6 IT Car3). Notably, L4/5 IT and L6 IT Car3 neurons are not provided in reference dataset, thus the predictions of all cell typing methods do not recall them. As shown in Fig. 2k, the extended experiment of Spatial-ID demonstrates its promising ability to discover these cell types. (**c**) Inhibitory neurons (Lamp5, Pvalb, Sncg, Sst and Vip).

**Extend Fig. 2.**
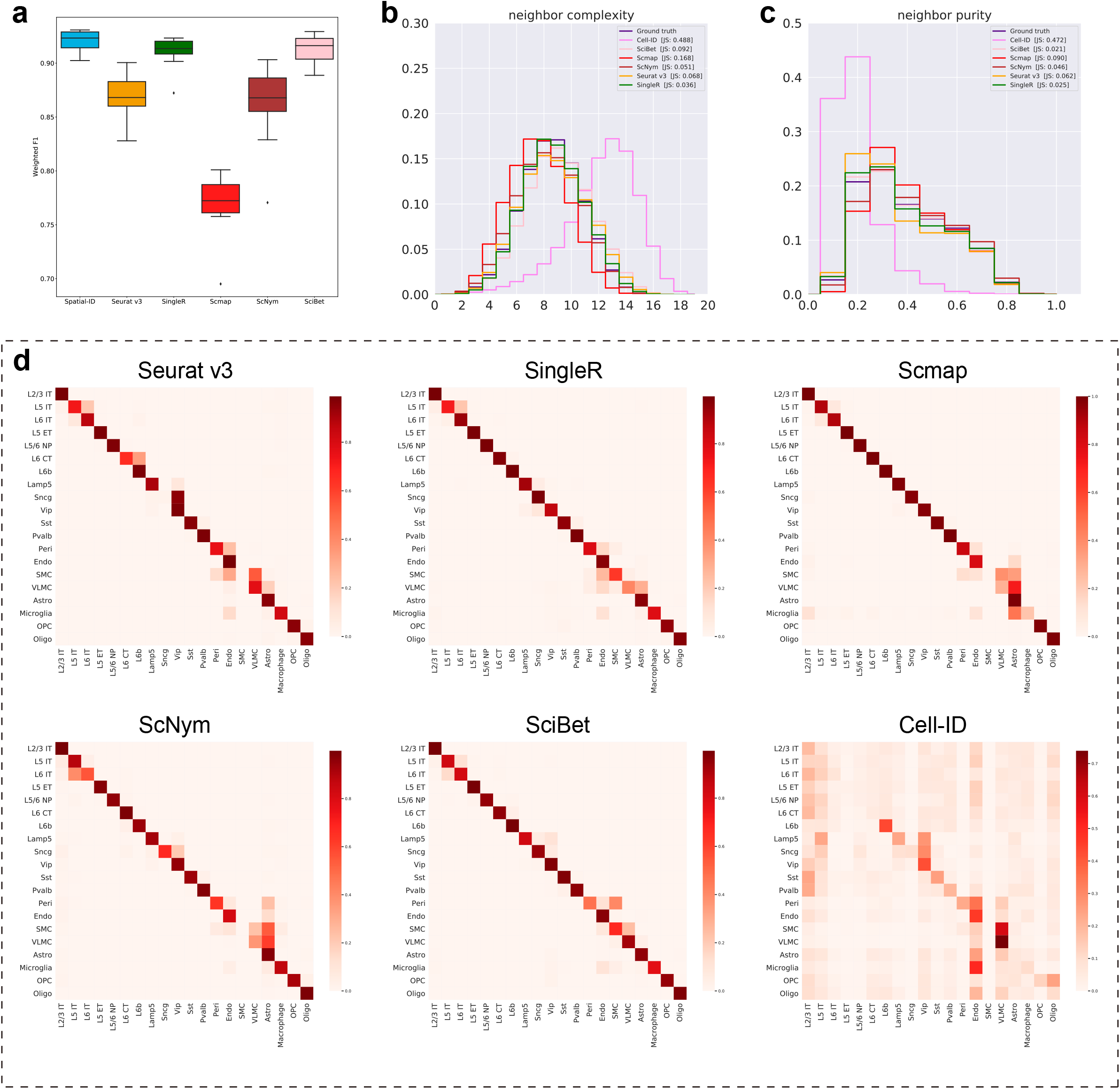
Additional comparisons for the mouse primary motor cortex dataset. **(a)** The comparison of mean weighted F1 score. The weighted F1 score of each sample is calculated by weighted averaging the F1 score of each cell type, in order to mitigate the effects of cell type imbalance. (**b**) Neighborhood complexity of the control methods. (**c**) Neighborhood purity of the control methods. (**d**). The confusion matrixes of the control methods.

**Extend Fig. 3.**
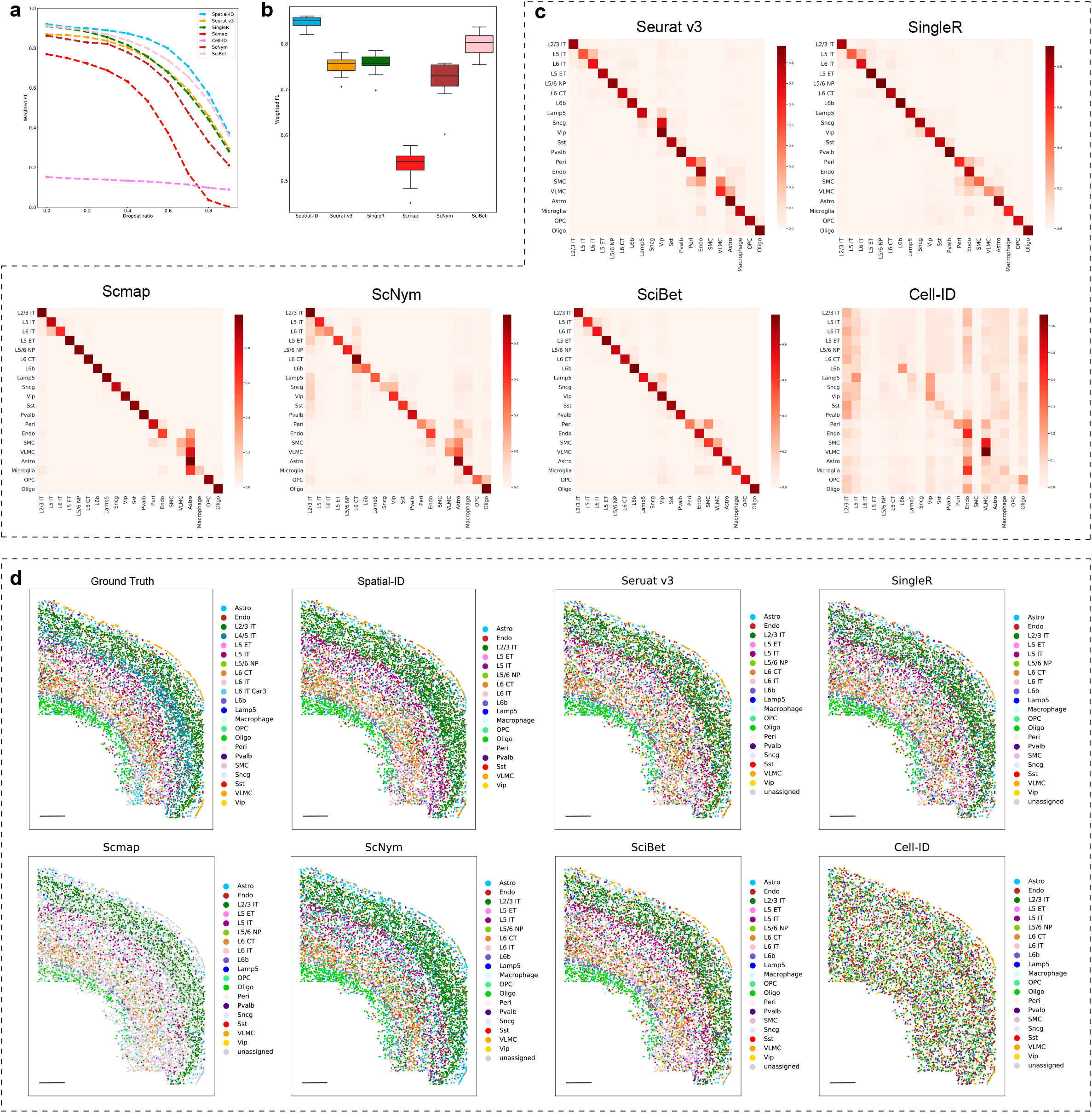
Neighborhood complexity of each cell type in mouse primary motor cortex dataset measured by MERFISH.

**Extend Fig. 4.**
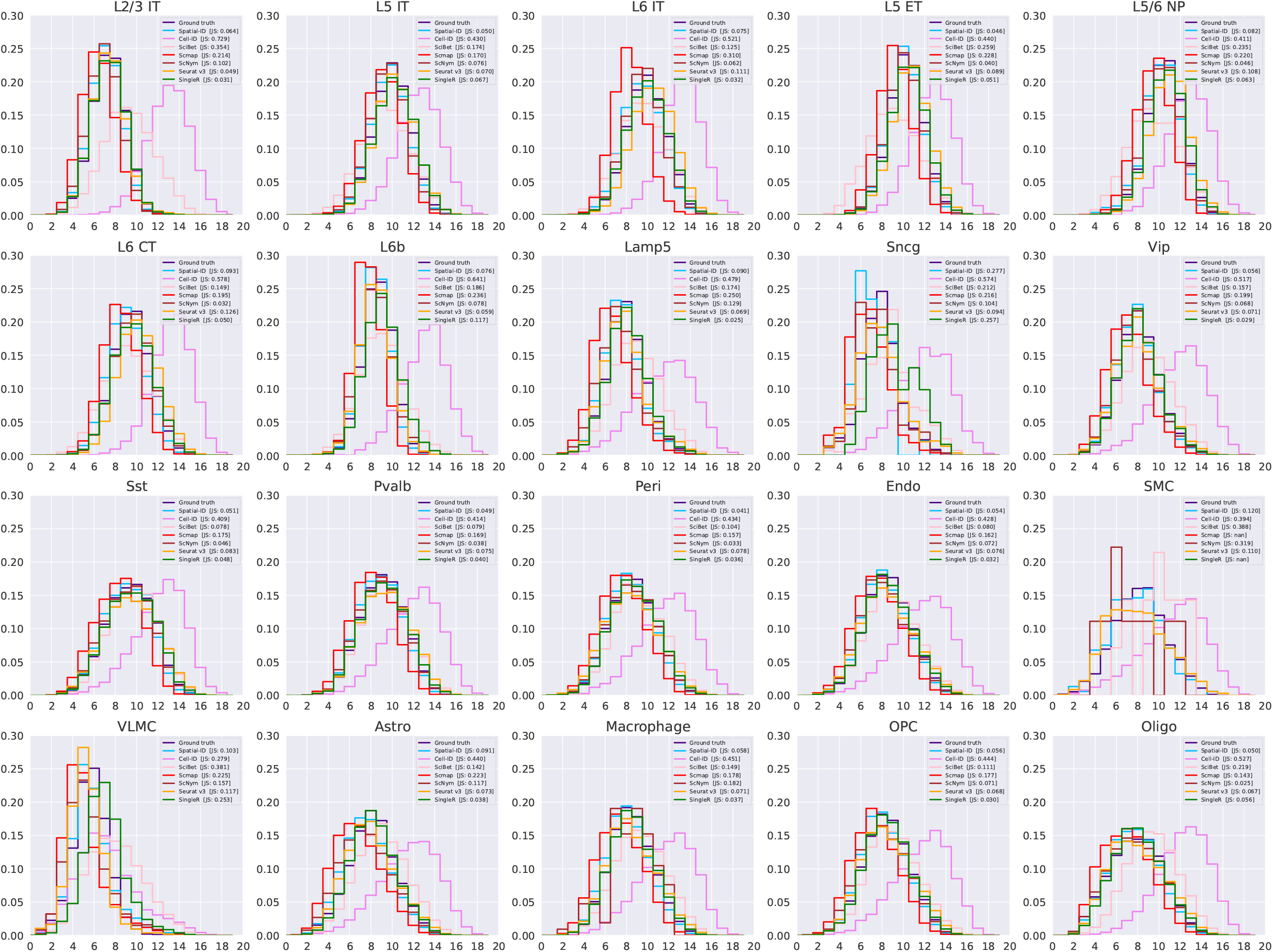
Neighborhood purity of each cell type in mouse primary motor cortex dataset measured by MERFISH.

**Extend Fig. 5.**
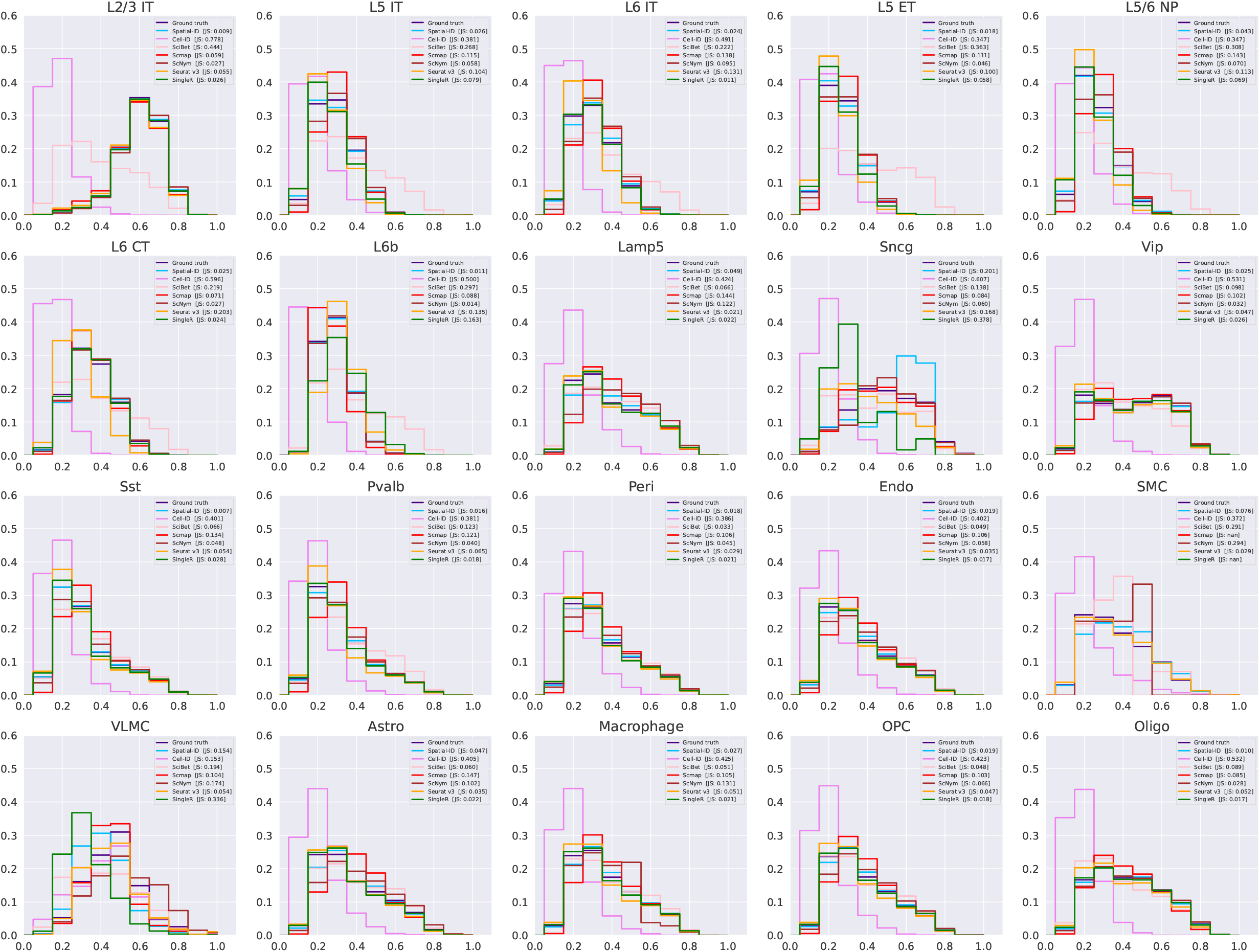
Additional comparisons of different gene dropout rates for the mouse primary motor cortex dataset. **(a)** The comparison of mean weighted F1 score at different gene dropout rates. **(b)** The comparison of mean weighted F1 score at 0.5 gene dropout rate. **(c)** The confusion matrixes of the control methods at 0.5 gene dropout rate. **(d)** Spatial organization of ground truth and predictions of different cell typing methods at 0.5 gene dropout rate.

**Extend Fig. 6.**
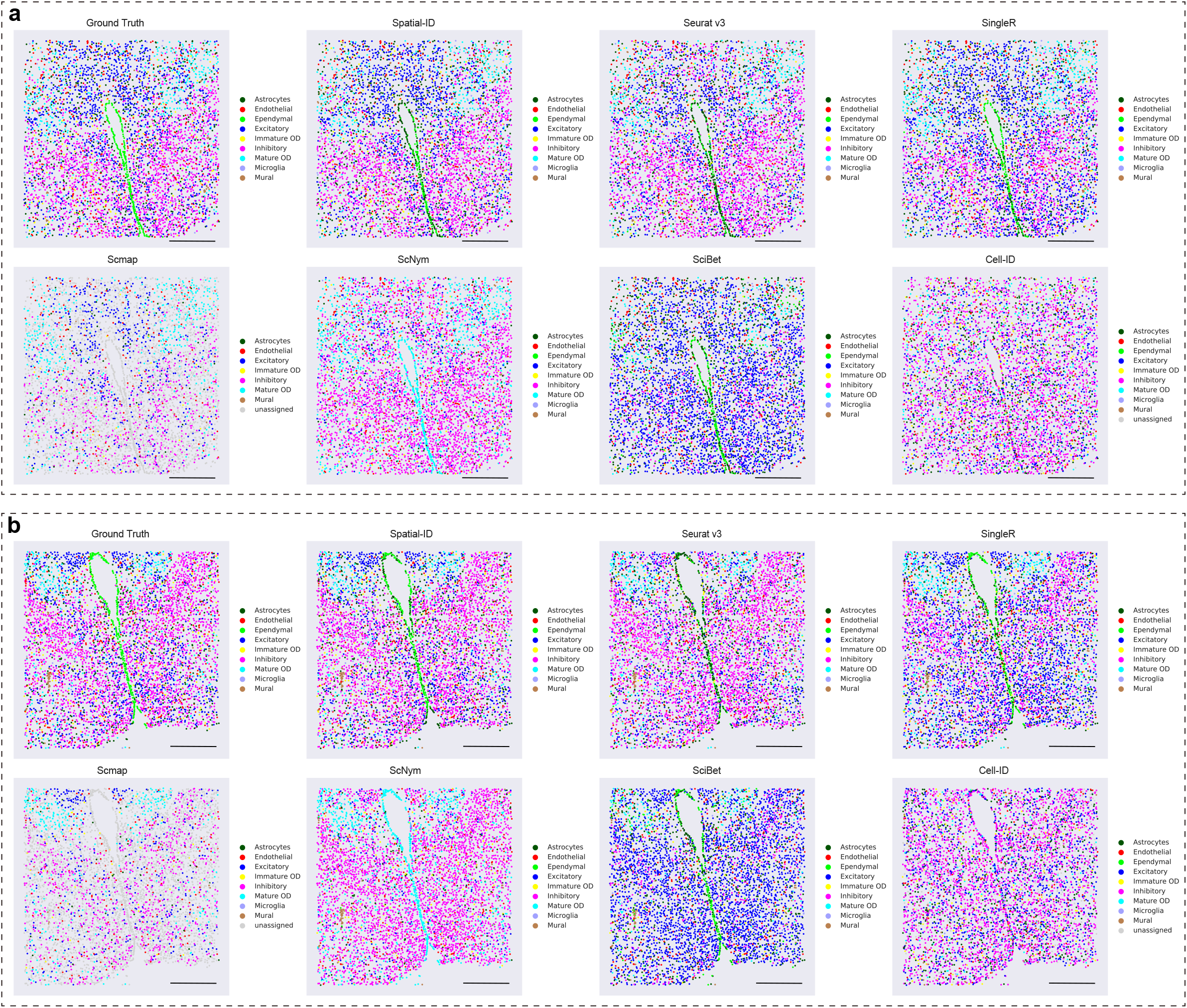
Additional comparisons of 2D spatial organization for the mouse hypothalamic preoptic region dataset. **(a)** The slice at Bregma −0.29. **(b)** The slice at Bregma −0.14.

**Extend Fig. 7.**
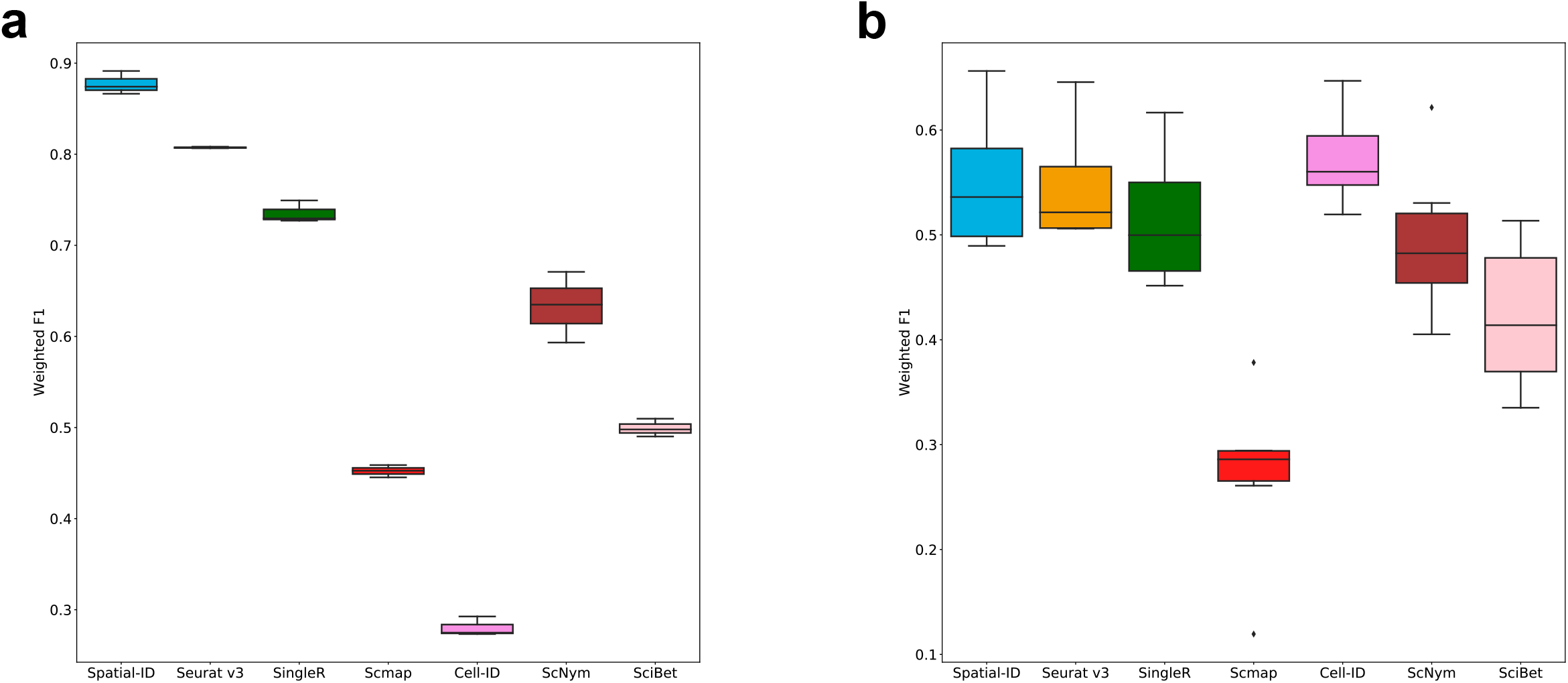
Additional comparisons of mean weighted F1 score. **(a)** The mouse hypothalamic preoptic region dataset measured by MERFISH. **(b)** The mouse spermatogenesis dataset measured by Slide-seq.

**Extend Fig. 8.**
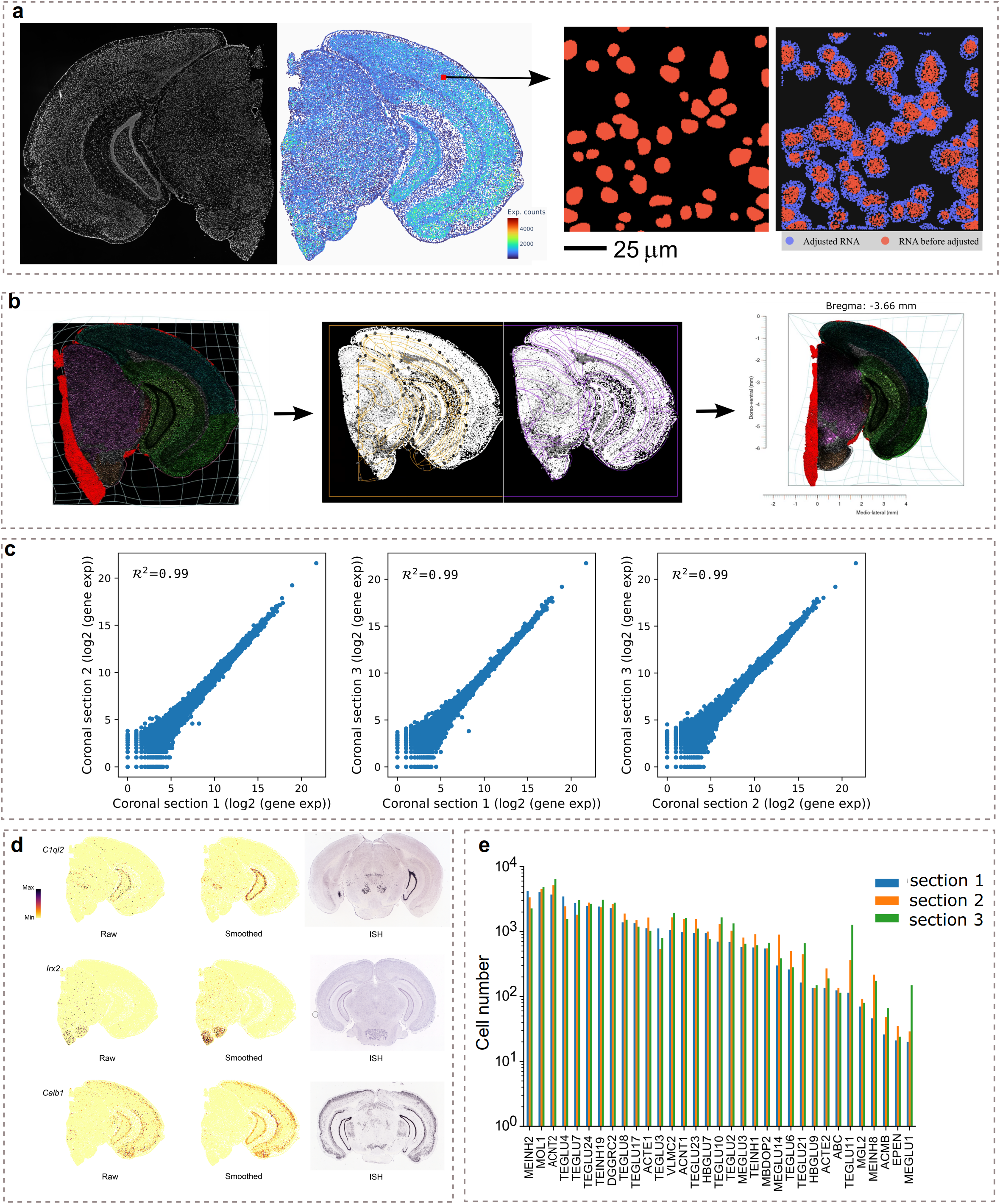
Data processing of the mouse brain hemisphere, and cell number distributions of identified cell types across 3 sections. (**a**) Cell segmentation procedure. ssDNA image of coronal section 3 and the corresponding spatial heatmap that indicates the number of transcripts captured by Stereo-seq are shown in the left panel. The right panel shows the segmented nuclei masks and transcripts surrounding the nucleus adjusted by a Gaussian mixture model. (**b**) The registration procedure is used to align coronal sections with the 3D Allen reference atlas. (**c**) Pearson’s correlation coefficients of spatial single-cell profiles between 3 adjacent coronal sections. (**d**) Comparisons between raw expression (left), smoothed by the Gaussian smoothing strategy (middle) and Allen ISH image (right) of selected genes *Clql2*, *Irx2* and *Calb1*. After Gaussian smoothing, these gene expressions show better spatial distributions that are much closer to their spatial captures of the in situ hybridization of Allen brain atlas. (**e**) Cell number of each identified cell type for the coronal sections 1, 2, and 3, respectively.

**Extend Fig. 9.**
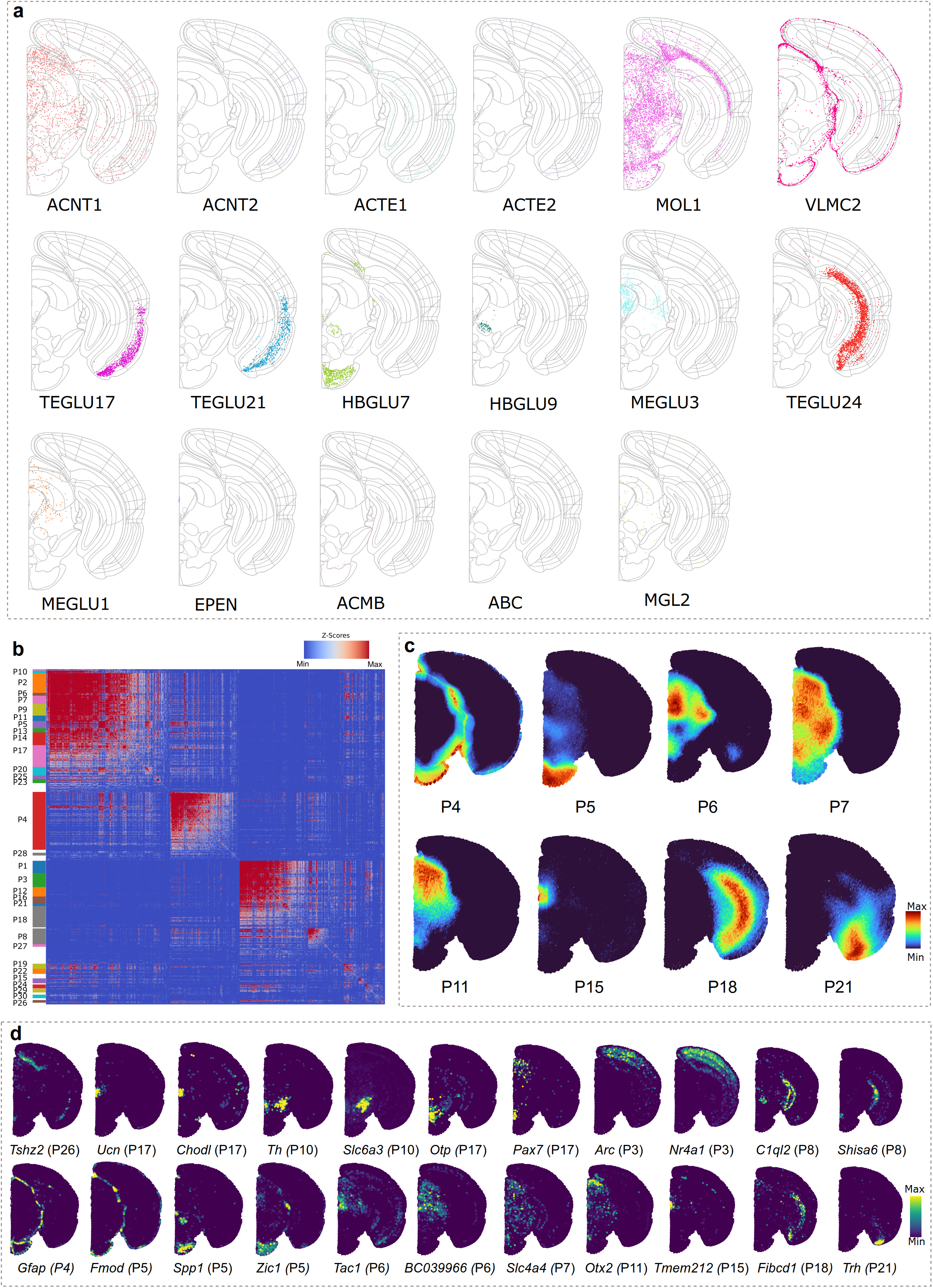
Additional spatial gene patterns and spatial heatmap of top scored genes. (**a**) Other identified cell types of Section 3. (**b**) Clustered spatial gene patterns. Genes with significant spatial correlation (FDR < 0.05) are clustered into 30 gene patterns on the basis of pairwise spatial correlation. (**c**) Some spatial gene patterns visualize with pattern scores. (**d**) Spatial heatmaps of top scored genes for corresponding spatial gene patterns indicate the number of transcripts captured by Stereo-seq.

## Notes

### Competing Interest Statement

The authors have declared no competing interest.

